# Compensatory role of KatG in defending H_2_O_2_ stress in *msr* deletion strain of *Salmonella* Typhimurium

**DOI:** 10.1101/2024.09.03.610964

**Authors:** Lalhmangaihzuali Lalhmangaihzuali, Suchitra Upreti, Raj Sahoo, Tapan Kumar Singh Chauhan, Manish Mahawar

## Abstract

KatG and Msrs are important enzymes associated with ROS homeostasis and bacterial survival under oxidative stress. Consistent to this notion, mutant strains in these enzymes showed hypersensitivity to oxidants and accumulates elevated levels of ROS. In current study we observed that a pan *msr* deletion (Δ5*msr* mutant) strain of *S.* Typhimurium accumulates significantly higher levels of ROS. However, unexpectedly, as compared to *S.* Typhimurium, the Δ5*msr* mutant strain exhibits more than 2000 folds resistance to H_2_O_2_. Transcriptional and mass spectrometry analyses reveal the upregulation of KatG in Δ5*msr* mutant strain. Further, Δ5*msr* mutant strain exhibits ∼6 folds higher KatG activity. Supplementation of Δ5*msr* mutant culture with reduced glutathione resulted in ROS neutralization, decreased KatG activity and abrogation of H_2_O_2_ resistance. However, Δ5*msr* mutant strain showed negligible KatE and KatN activities. The findings of current study suggest that the *Salmonella* have evolved the mechanism to upregulate one antioxidant gene in absence of others to mitigate oxidative stress.

## Introduction

Nontyphoidal *Salmonella* (NTS) serovars are the major cause of gastroenteritis. NTS serovars are responsible for approximately 150 million cases of illness and about 60,000 annual deaths worldwide (CDC, 2024). *Salmonella enterica* serovar Typhimurium (*S.* Typhimurium) and *Salmonella enterica* serovar Enteritidis (*S.* Enteritidis) are the most prevalent NTS serovars (Gilchrist and MacLennan, 2019). Multidrug resistance and resistance to third-generation cephalosporins, such as ceftriaxone (Balasubramanian *et al*., 2019) led NTS to be listed as high priority organism.

*S.* Typhimurium suffers but survives oxidative stress. *S.* Typhimurium encounters reactive oxygen species (ROS) from two key sources. During infection process, the ROS is generated by host inflammatory cells. By sensing bacterial ligands, phagocytic cells assemble NADPH oxidase and generate copious amounts of superoxides (O_2_ ^-^). Other very important source of ROS in *S*. Typhimurium is its own metabolism where superoxides are generated by incomplete reduction of molecular O_2_ and by the autoxidation of enzymes like dehydrogenases, glutathione reductase, and cytochromes P450 (Imlay, 2013; Seixas *et al*., 2022). O_2_^−^ has a half-life in the range of microseconds, however, it can oxidize several targets like ascorbate and thiols (Kehrer, 2000). H_2_O_2_ is produced by the dismutation of O_2_^−^ and autoxidation of flavoenzymes (Imlay, 2013). H_2_O_2_ with a half-life in the range of minutes, is one of the most important and stable oxidants (Kehrer, 2000). Further, H_2_O_2_ interacts with metal cofactors such as Fe^2+^ and generates the highly reactive OH^•^ radicals (Fasnacht and Polacek, 2021). ROS can damage bacterial DNA, RNA, lipids and proteins (Fasnacht and Polacek, 2021; Seixas *et al*., 2022) leading to cell death. Proteins, due to abundance and reactive nature, are highly susceptible to the ROS mediated damage. The ROS can oxidize almost all amino acids, however, Met residues, because of the presence of sulfur atom are highly vulnerable to oxidation, leading to the formation of methionine sulfoxide (Met-SO) (Levine *et al*., 1996).

To survive under oxidative stress, the bacterial pathogens, including *S.* Typhimurium have evolved various primary antioxidants and repair enzymes. *S.* Typhimurium mitigates ROS mediated damage directly by degrading ROS and by repairing the oxidized amino acid residues. *S.* Typhimurium employs four superoxide dismutases (Sods, SodA, SodB, SodCI, and SodCII) (Fang *et al*., 1999; Imlay, 2013) which catalytically degrades O_2_^-^. *S.* Typhimurium encodes three catalases (KatE, KatG, and KatN), and three peroxiredoxins (AhpC, TsaA, and Tpx) which help in neutralising H_2_O_2_ (Robbe-Saule *et al*., 2001; Hebrard *et al*., 2009; Imlay, 2013; Kroger *et al*., 2013). While AhpC scavenges low concentrations (μM) of H_2_O_2_, catalases are the more efficient in degrading high concentrations (mM) of H_2_O_2_ (Hillar *et al*., 2000; Mishra and Imlay, 2012). Among three catalases, the expression of KatG (also known as hydroperoxidase I, HPI) predominates during the exponential growth of *S.* Typhimurium (Kroger *et al*., 2013). KatG is known to contribute to the growth of *S.* Typhimurium under H_2_O_2_ stress *in vitro* and survival *in vivo* (Hebrard *et al*., 2009; Hahn *et al*., 2021; Kirthika *et al*., 2022). The expression of KatG is shown to be highly upregulated during oxidative stress in *S.* Typhimurium (McLean *et al*., 2010; Kroger *et al*., 2013; Kirthika *et al*., 2022). On other hand, the deletion of *katG* gene has been reported to enhanced accumulation ROS in *E. coli* (Luan *et al*., 2018) and *S.* Typhimurium (Kirthika *et al*., 2022).

By repairing Met-SO to Met, methionine sulfoxide reductases (Msrs) provides a mean to quench ROS thus enhances the cellular survival under oxidative stress conditions (Mahawar *et al*., 2011; Benoit *et al*., 2013; Kuhns *et al*., 2013; Gennaris *et al*., 2015; Sarkhel *et al*., 2017; Andrieu *et al*., 2023). *S.* Typhimurium encodes five Msrs: four cytoplasmic MsrA, MsrB, MsrC, and BisC, and one periplasmic MsrP. While MsrA repairs both protein-bound and free Met-*S*-SO, MsrB targets protein-bound Met-*R*-SO. MsrC repairs free Met-*R*- SO, and BisC tackles biotin sulfoxides as well as free Met-*S*-SO (Denkel *et al*., 2011; Denkel *et al*., 2013; Trivedi *et al*., 2015; Nair *et al*., 2021). MsrP repairs both protein-bound and free Met-*R*-SO and Met-*S*-SO (Andrieu *et al*., 2020; Shome *et al*., 2020; Chandra *et al*., 2023). Cytoplasmic Msrs mediated repair requires TrxA and TrxR where NADPH serves as an electron donor (Dixit *et al*., 2017). However, MsrP mediated repair uses electrons from respiratory chain which are channel through MsrQ (Ezraty *et al*., 2017).

Msr-mediated repair serves two key functions in the cell: first, it restores the functions of Met-SO containing proteins (Ezraty *et al*., 2004; Khor *et al*., 2004; Mahawar *et al*., 2011; Benoit *et al*., 2013; Kuhns *et al*., 2013; Sarkhel *et al*., 2017; Nasreen *et al*., 2022). In fact, proteins with higher percentage of Met residues tend to resist loss of function under oxidizing conditions in comparison to other amino acid (Levine *et al*., 1999). Second, it maintains ROS homeostasis in the cell (Abulimiti *et al*., 2003; Luo and Levine, 2009; Benoit and Maier, 2016; Schmalstig *et al*., 2018). The Met residues, either in free or in protein bound form get oxidized and convert in to Met-SO thereby quench excess oxidants and prevent macromolecular damage. Later, these oxidized Met-SOs are reduced by Msr machinery providing a mechanism to sop up excess oxidants (Stadtman *et al*., 2003).

We hypothesized that a pan *msr* strain of *S*. Typhimurium should show perturbed ROS homeostasis and hypersensitivity to ROS. Indeed, earlier we have observed very high susceptibility of Δ5*msr* mutant strain to oxidants like HOCl, Chloramine-T and paraquat (Sahoo *et al*., 2023). In contrast to above observation, the results of current study reveal very high resistance of Δ5*msr* mutant strain to H_2_O_2_. Further, we show the accumulation of ROS and very high KatG activity in Δ5*msr* mutant strain.

## Results

### Δ5*msr* mutant strain shows very high resistance to hydrogen peroxide

H_2_O_2_ is one of the key oxidants generated by the phagocytic cells (Mastroeni *et al*., 2000) and Met residues are the prime targets of H_2_O_2_ mediated oxidation (Luo and Levine, 2009; Kuhns *et al*., 2013). *msr* gene deletion strains of various bacterial pathogens showed hypersensitivity to H_2_O_2_ (St. John *et al*., 2001; Alamuri and Maier, 2004; Vattanaviboon *et al*., 2005; Atack and Kelly, 2009; Lei *et al*., 2011; Jalal and Lee, 2020). We evaluated the sensitivity of Δ5*msr* mutant strain of *S.* Typhimurium to H_2_O_2_. Unexpectedly, in comparison to *S.* Typhimurium, Δ5*msr* mutant strain has been more than 2000 folds more resistance to H_2_O_2_. The observed numbers (log_10_ CFUs; mean ± S.E.M.) following incubation of *S.* Typhimurium and Δ5*msr* mutant strains with 25 mM H_2_O_2_ were 5.85 ± 0.32 and 9.2 ± 0.046 respectively. Following incubation with 50 mM H_2_O_2_, we did not recover any viable bacteria in case of *S.* Typhimurium, however, recovered numbers (log_10_ CFUs; mean ± S.E.M.) in case of Δ5*msr* mutant strain were 8.91 ± 0.035 (Fig. 1).

**Fig. 1:**
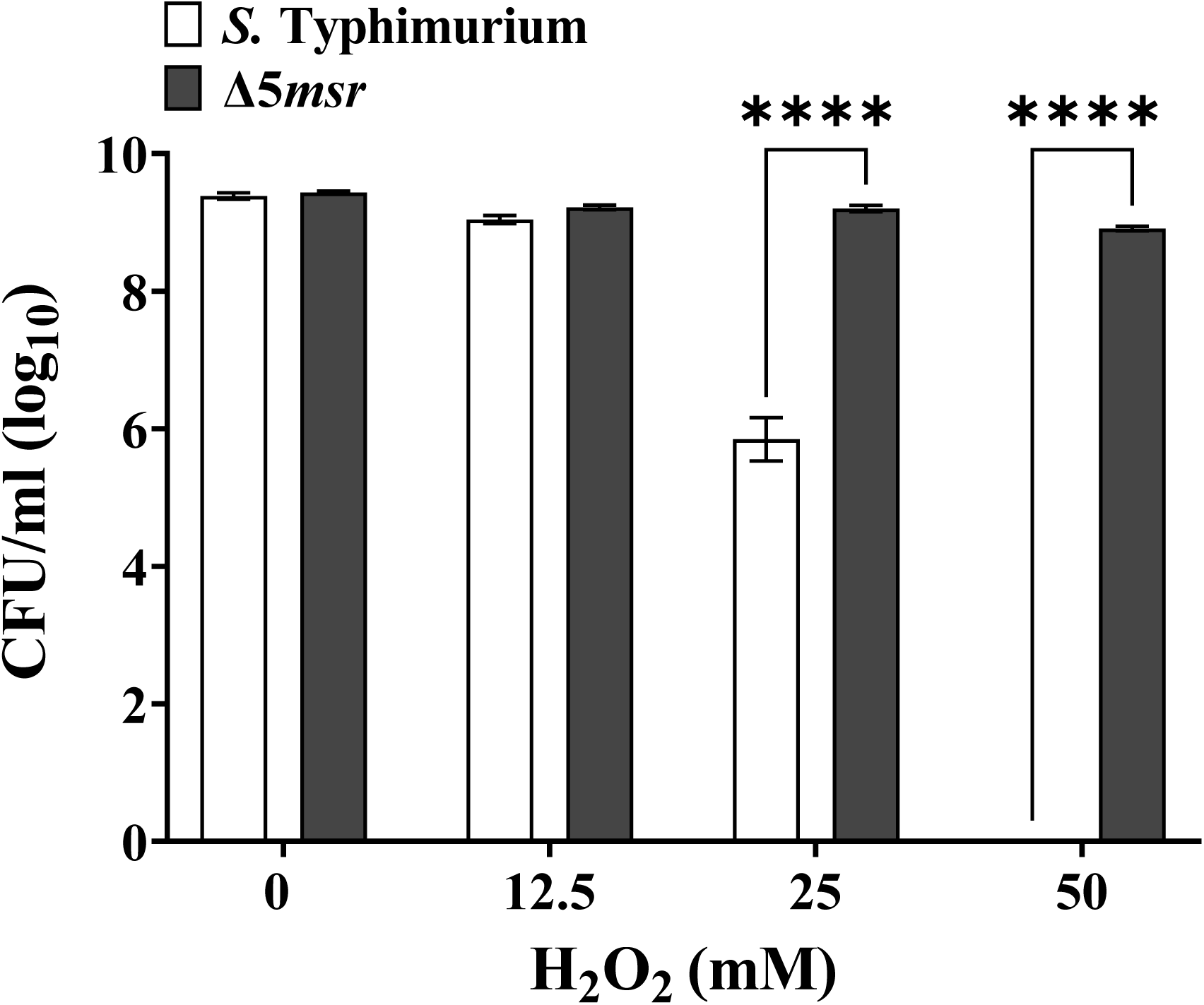
Δ5*msr* mutant strain shows high resistance to H_2_O_2_. Mid log grown cultures of *S.* Typhimurium and Δ5*msr* mutant strains were exposed to H_2_O_2_ for 2 hours. The cultures were then serially diluted and plated on HE agar plates. The CFUs were counted following overnight incubation of the plates. Data are presented as mean ± S.E.M. (*n* = 6). \*\*\*\**p* < 0.0001

### Δ5*msr* mutant strain exhibits very high catalase specific activity

Catalase and AhpC are the key H_2_O_2_ degrading enzymes (Hillar *et al*., 2000; Mishra and Imlay, 2012). Exposure of *S.* Typhimurium to H_2_O_2_ has been shown to enhance the expression of heme catalases (Hahn *et al*., 2021). Next, we estimated the catalase specific activity in the cell free lysates of *S.* Typhimurium and Δ5*msr* mutant strains. We observed more than 15 folds higher (*p* < 0.0001) catalase specific activity in Δ5*msr* mutant strain as compared to *S*. Typhimurium (Fig. 2a).

**Fig. 2:**
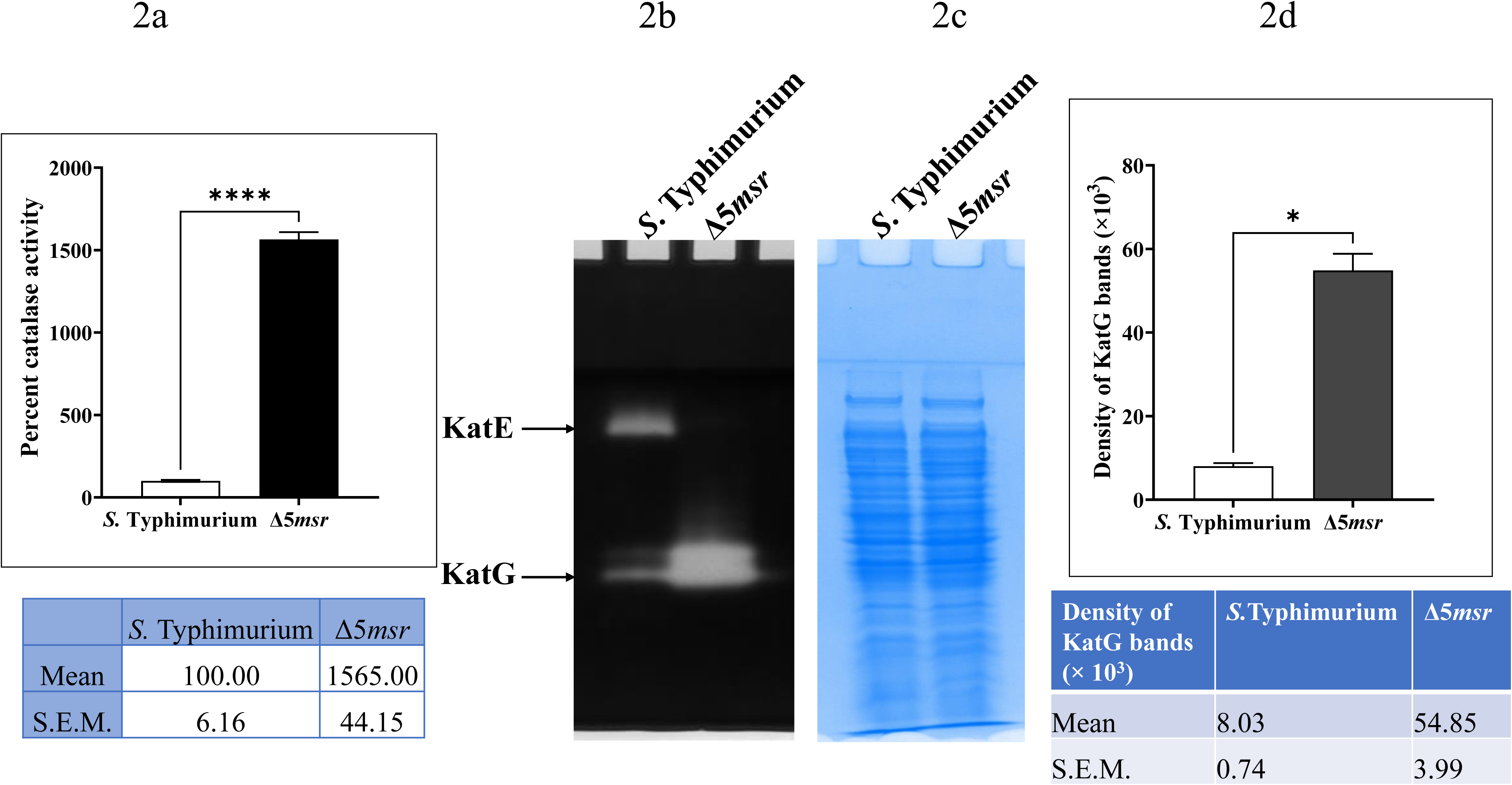
Δ5*msr* mutant strain exhibits significantly higher KatG activity. Mid log grown cultures of *S.* Typhimurium and Δ5*msr* mutant strains were pelleted and suspended in PBS and lysed by sonication. Catalase specific activity was determined as detailed in materials and methods (Fig. 2a). Activity in *S.* Typhimurium was considered as 100%. Data are presented as mean ± S.E.M. (*n* = 8). \*\*\*\**p* < 0.0001 For activity estimation on native gel, 50 µg cell free lysates of *S.* Typhimurium and Δ5*msr* mutant strains were resolved on native gel and the catalase activity was determined as described in the material and methods (Fig. 2b). The duplicate native gel stained with CBB (Fig. 2c) served as a loading control. The densitometric analysis using ImageJ software is depicted in Fig. 2d. Data are presented as mean ± S.E.M. (*n* = 2) \**p* < 0.05

### Δ5*msr* mutant strain depicts significantly higher KatG activity

*S.* Typhimurium encodes three catalases viz. KatE, KatG and KatN (Robbe-Saule *et al*., 2001). We analysed the catalase activity on native gel (Weydert and Cullen, 2010). Interestingly, Δ5*msr* mutant strain showed more than 6 folds higher KatG activity as compared to *S.* Typhimurium (Fig. 2b, 2d). In these assays, the gels stained with CBB served as a loading control (Fig. 2c). In contrast, we observed very low KatE activity in vigorously growing cells which further increased in late log to stationary phase cultures of *S.* Typhimurium strain. However, KatN activity was detectable in very late stationary phase cultures of *S.* Typhimurium. Interestingly, no perceivable activities of KatE and KatN were detected in the Δ5*msr* mutant strain (Fig 3a). In these assays, the gels stained with CBB served as a loading control (Fig. 3b).

**Fig. 3:**
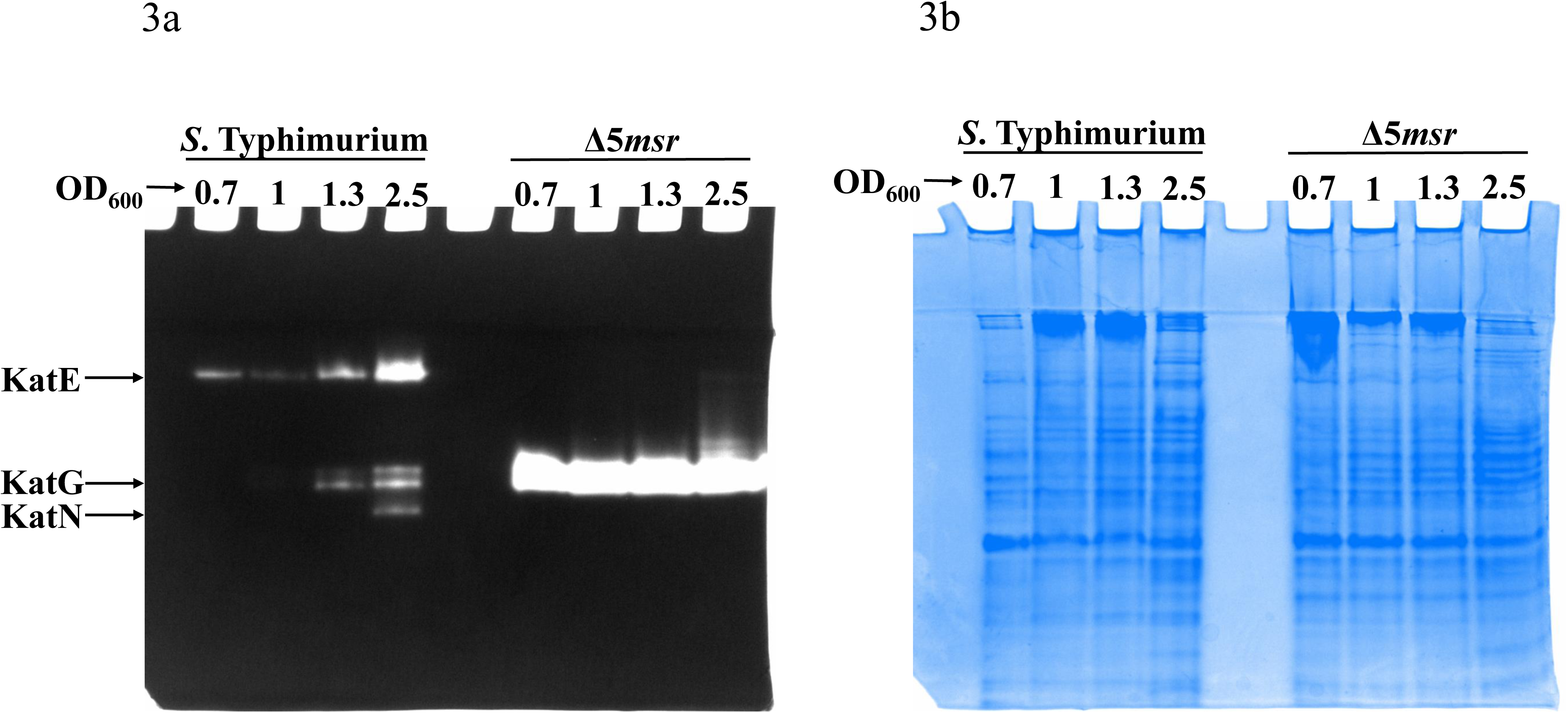
Differential expression of various catalases during different growth phases of *S.* Typhimurium and Δ5*msr* mutant strains. *S.* Typhimurium and Δ5*msr* mutant strains were grown in LB broth at indicated OD_600_. 50 µg of cell free lysates from such cultures were resolved on native gel and the catalase activity was determined as described in the material and methods (Fig. 3a). Duplicate native gel followed by CBB staining was used as loading control (Fig. 3b).

### Mass spectrometry and transcriptional analyses reveal upregulation of KatG in Δ5*msr* mutant strain

SDS-gel analysis revealed an upregulated band of ∼80 kDa in Δ5*msr* mutant strain (Fig. 4a). Mass spectrometry based analysis identified upregulated band as KatG (Supplementary Fig. S1). Further, RT-qPCR analysis reveals more than 170 folds higher *kat*G transcription in the Δ5*msr* mutant strain as compared to *S.* Typhimurium (Fig. 4b). In contrast, RT-qPCR based analysis shows 2.89 and 2.06 folds lower transcription of *katE* (Fig. 4c) and *katN* (Fig. 4d) in the Δ5*msr* mutant strain as compared to *S.* Typhimurium.

**Fig. 4:**
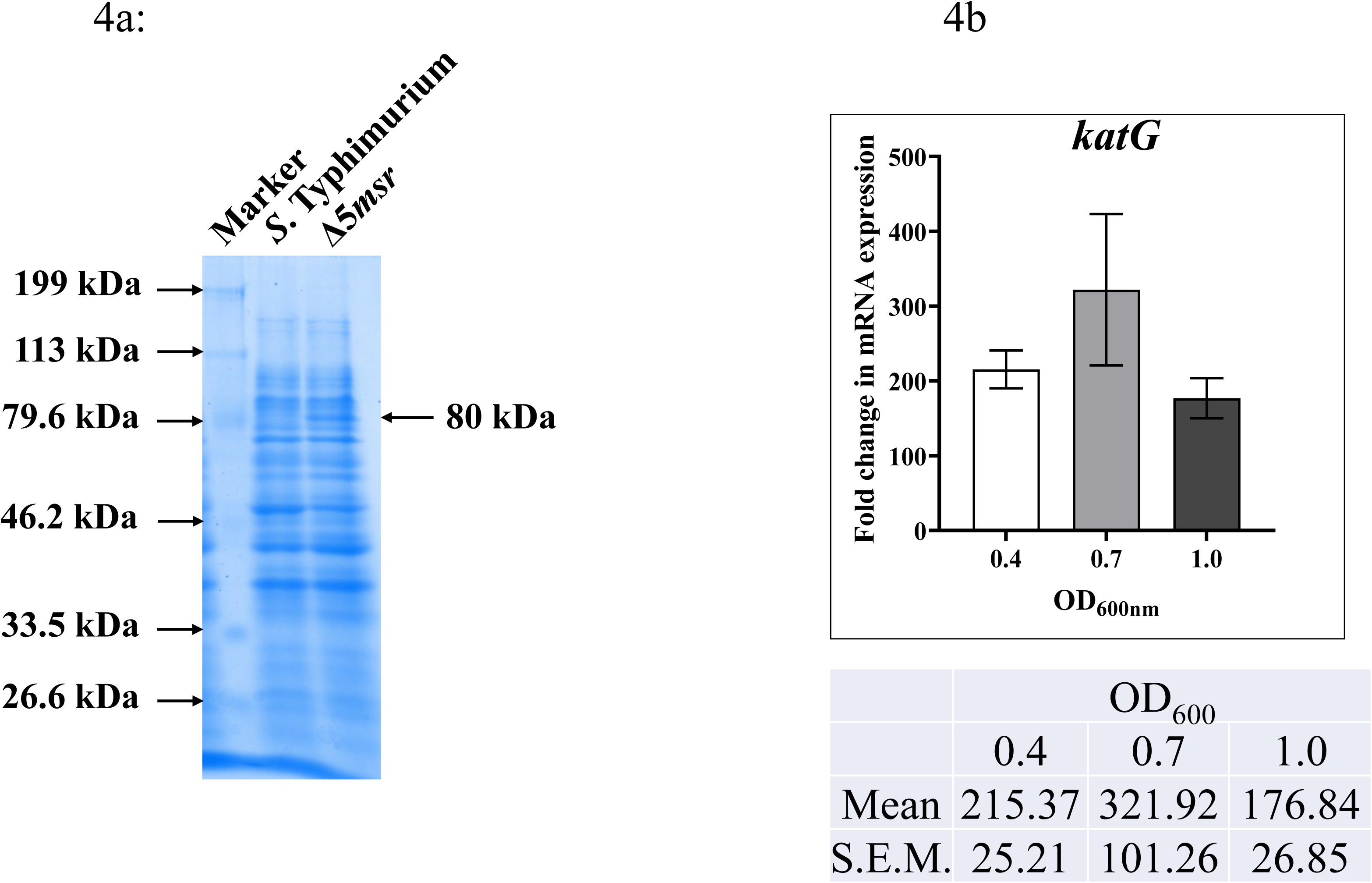
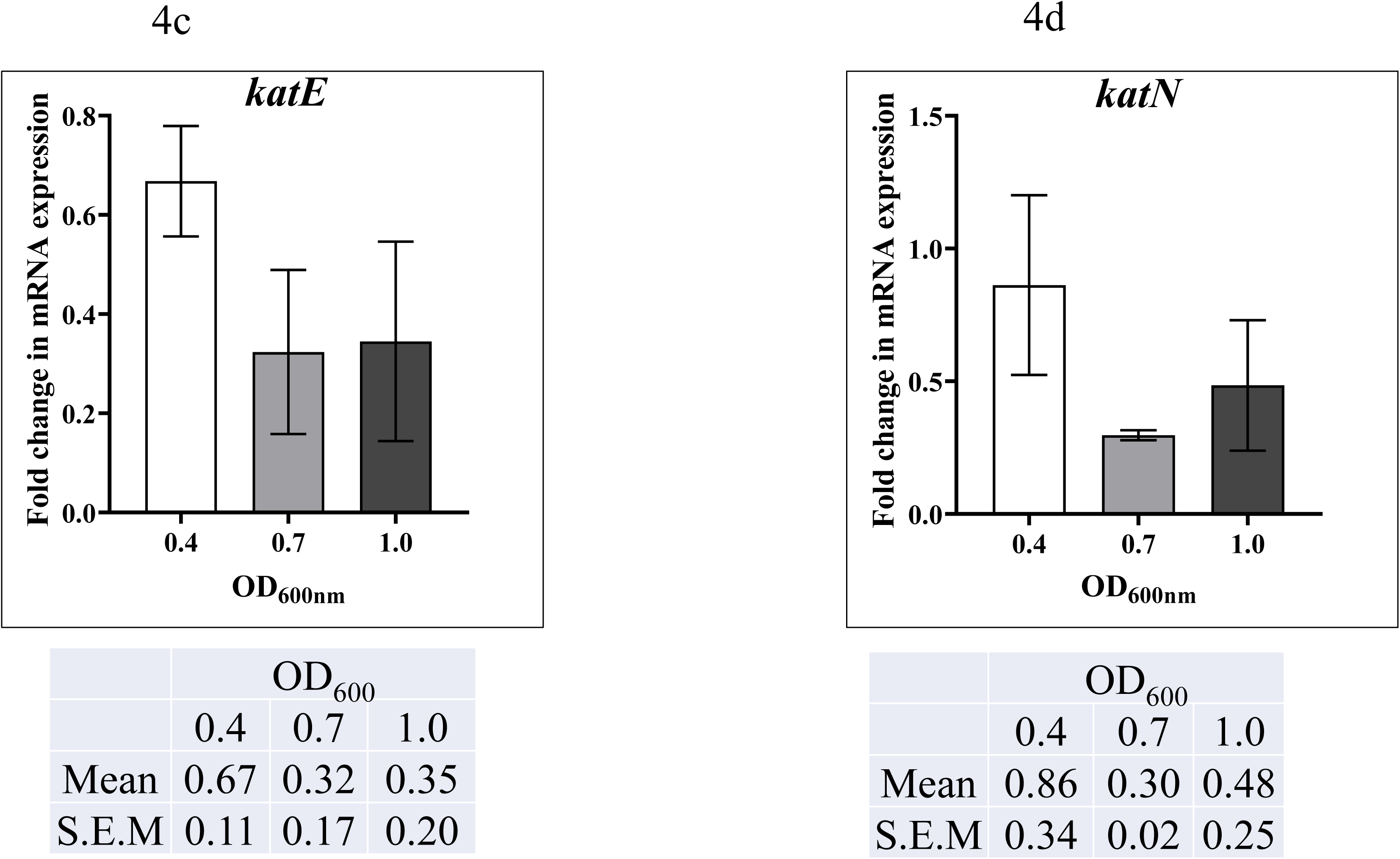
Mass spectrometry and transcriptional analysis reveal upregulation of KatG in Δ5*msr* mutant strain. Lysates of Mid-log grown cultures of *S*. Typhimurium and Δ5*msr* mutant strains were resolved in 10% SDS gel and stained with CBB. The upregulated protein band in Δ5*msr* mutant strain was incised and identified by mass spectrometry (Fig. 4a). For RT-qPCR analysis, the cultures grown at various ODs were pelleted. RNA isolation and cDNA synthesis and qPCR were done as described in materials and methods. The relative expression levels (Δ5*msr* mutant strain to *S.* Typhimurium) of *katG* (Fig. 4b), *katE* (Fig. 4c) and *katN* (Fig. 4d) were calculated using the 2^-ΔΔCT^ method, with *gmk* serving as the housekeeping gene. The data are presented as mean ± S.E.M. (n = 4).

### Δ5*msr* mutant strain exhibits higher ROS but lower thiol levels

Msrs are known to maintain ROS homeostasis of cell (Abulimiti *et al*., 2003; Luo and Levine, 2009; Benoit and Maier, 2016; Schmalstig *et al*., 2018). Next, we determined the ROS levels in *S.* Typhimurium and Δ5*msr* mutant strains (Chandra *et al*., 2024). The ROS levels in *S.* Typhimurium were considered as 100%. We observed more than 5 folds increase in ROS levels in Δ5*msr* mutant strain as compared to *S.* Typhimurium strain (*p* < 0.01) (Fig. 5a). Fluorescence microscopy analysis reveals higher green fluorescence in Δ5*msr* mutant strain as compared to *S.* Typhimurium strain depicting higher levels of ROS (Fig. 5b). Conversely, Δ5*msr* mutant strain shows 1.4 folds lower total thiol levels than *S.* Typhimurium (Fig. 6).

**Fig. 5:**
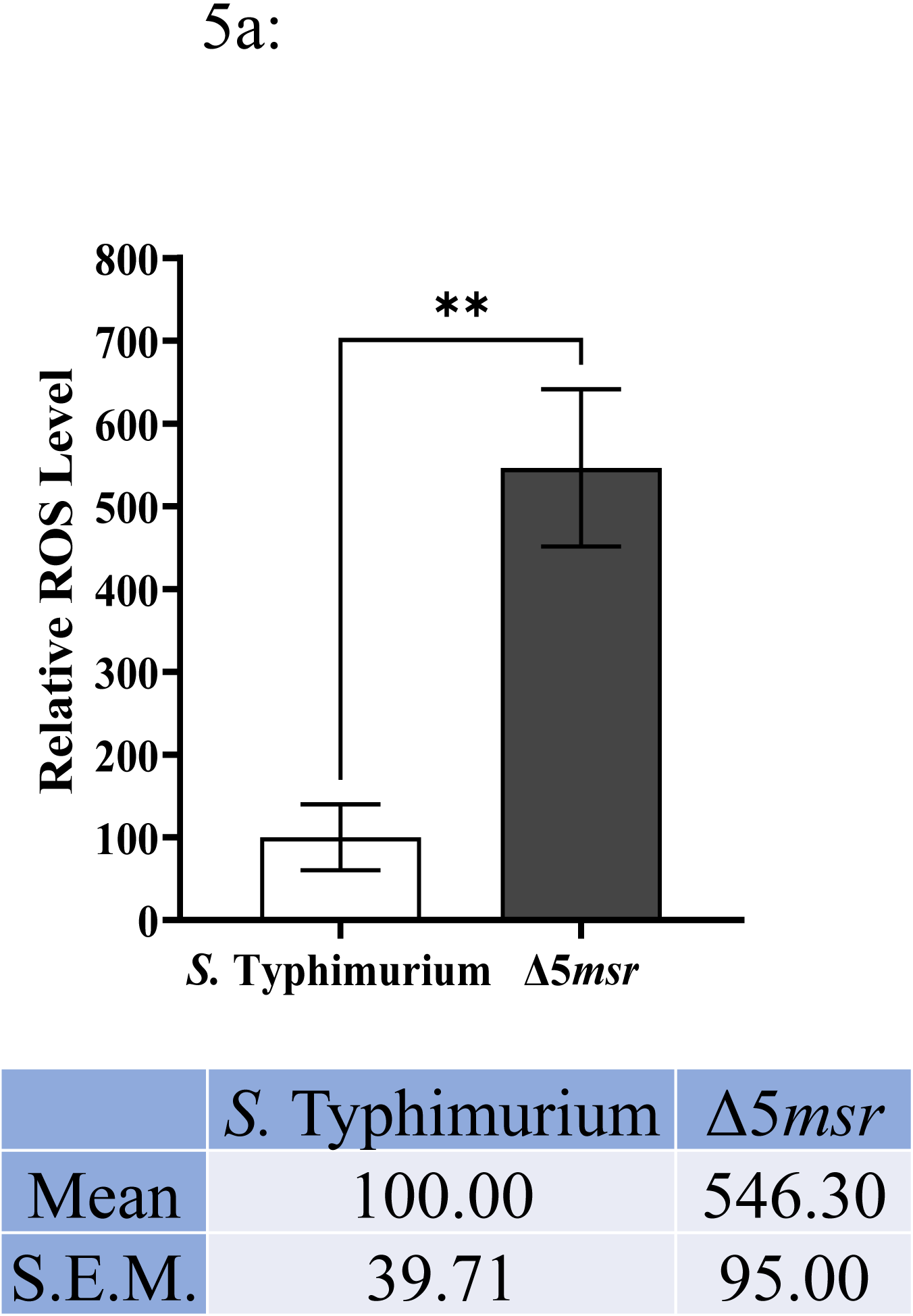
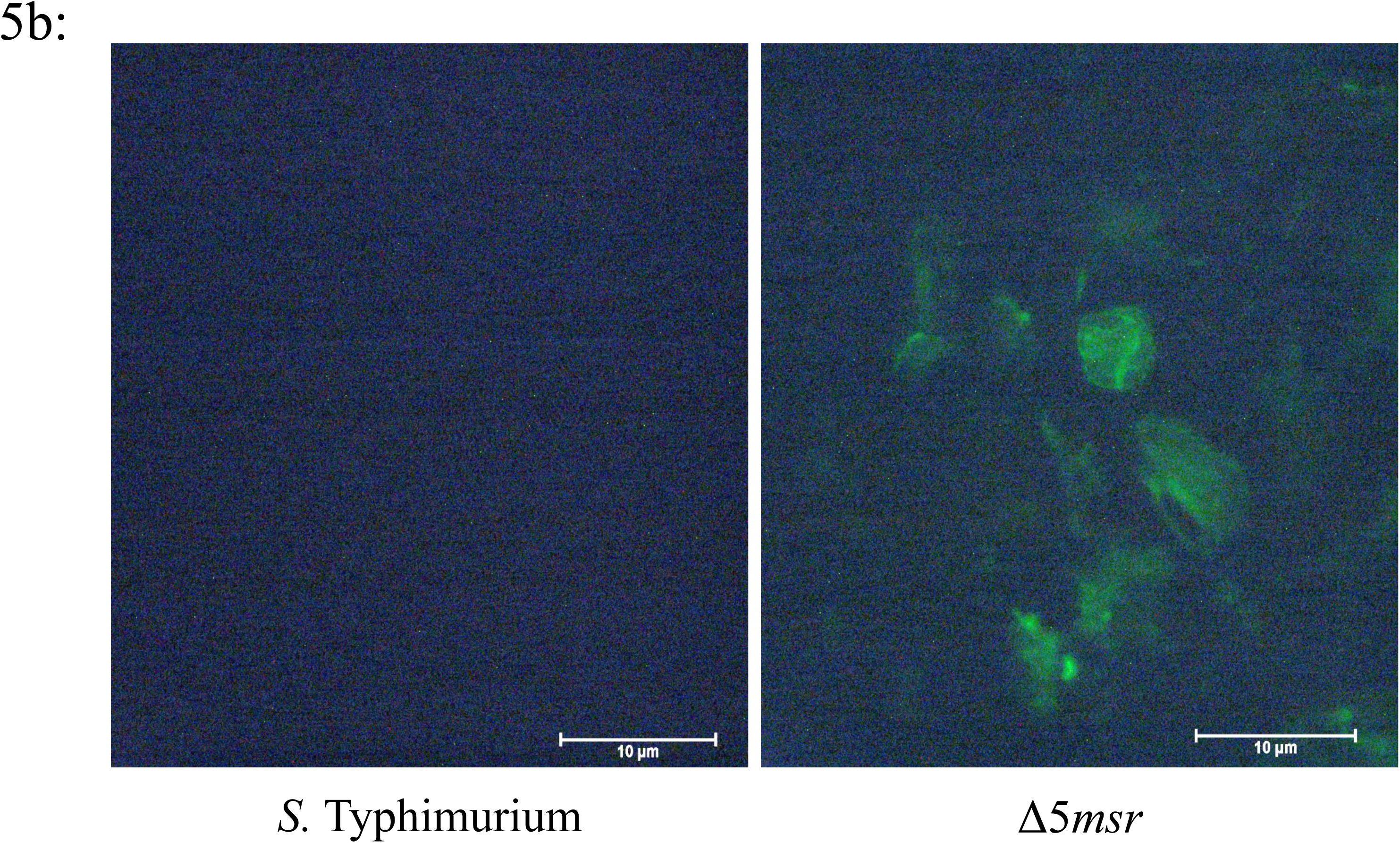
Δ5*msr* mutant strain accumulates higher levels of ROS. Vigorously growing cultures of *S.* Typhimurium and Δ5*msr* mutant strains were incubated with DCFDA. Following washing, the fluorescence was measured by fluorometer using excitation and emission at 485 and 535 nm, respectively (Fig. 5a) or observed under a fluorescent microscope (Fig. 5b). ROS levels of *S.* Typhimurium strain was considered as 100%. Data are presented as mean ± S.E.M. (*n* = 5). \*\**p* < 0.01

**Fig. 6:**
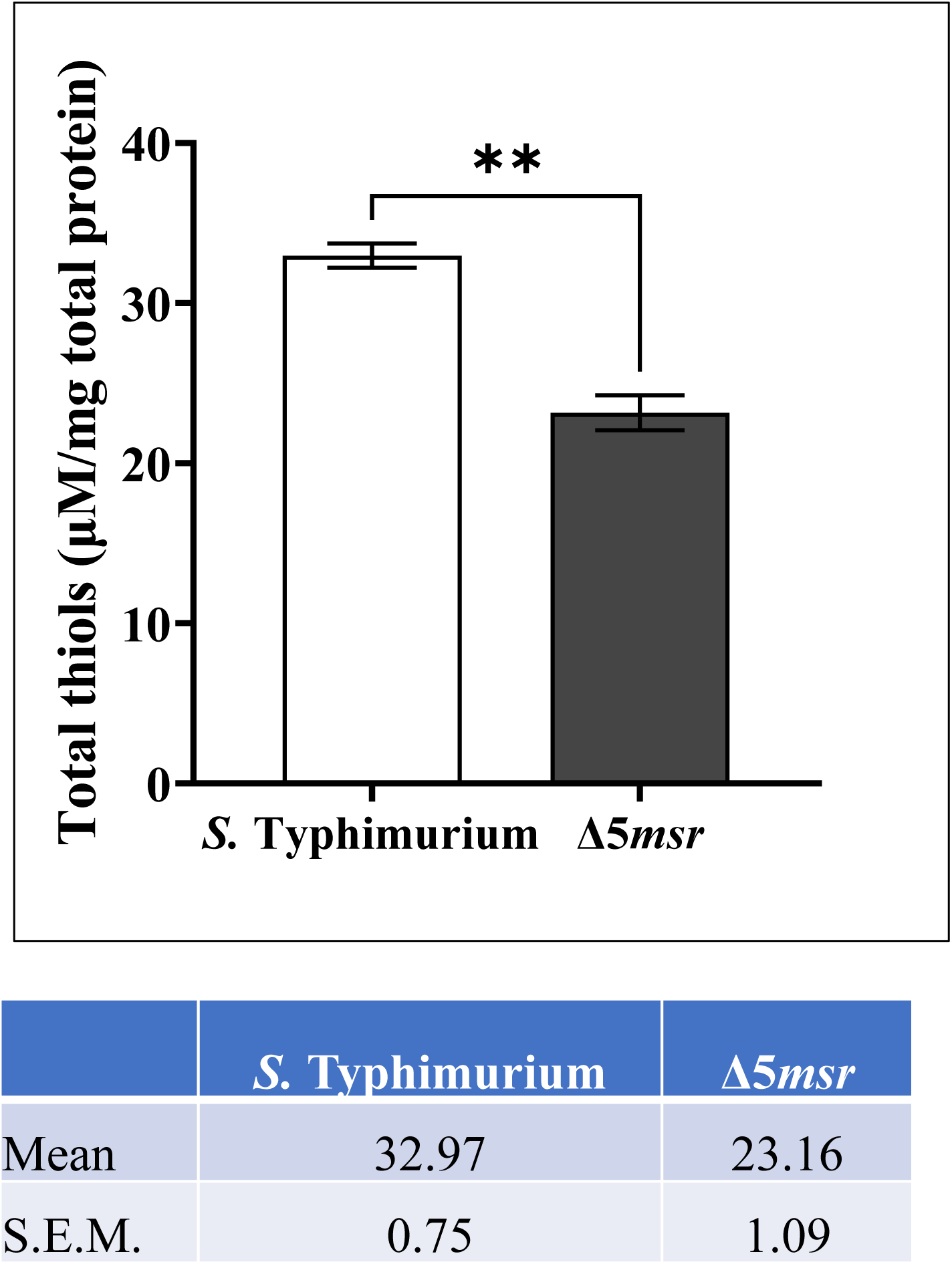
Δ5*msr* mutant strain shows lower total thiols. Total thiol levels in *S.* Typhimurium and Δ5*msr* mutant strain were determined as detailed in the materials and methods. Data are presented as mean ± S.E.M. (*n* = 6). \*\**p* < 0.01

### Supplementation of Δ5*msr* mutant strain with glutathione (GSH) reduces ROS levels and KatG activity

The ROS levels in *S.* Typhimurium and Δ5*msr* mutant strains after GSH supplementation were determined (Kwon *et al*., 2019). We observed 2.2 folds decrease in ROS levels in Δ5*msr* mutant strain following supplementation with GSH (Fig. 7).

**Fig. 7:**
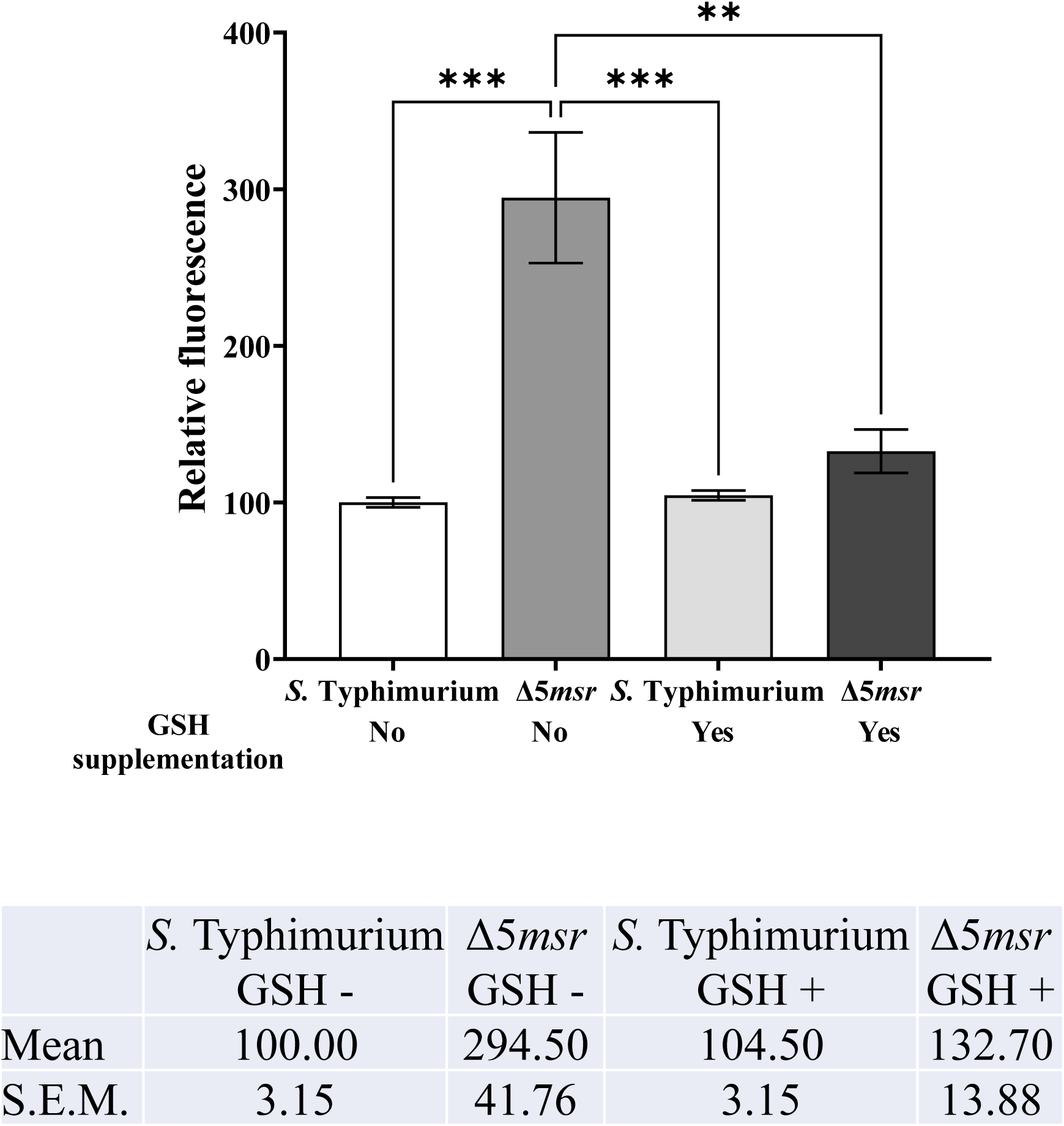
Supplementation of GSH reduces ROS levels in Δ5*msr* mutant strain. Vigorously growing cultures of *S.* Typhimurium and Δ5*msr* mutant strains (without supplementation or pre- supplemented with 10 mM GSH) were incubated with DCFDA. Fluorescence was measured by fluorometer using excitation and emission at 485 nm and 535 nm, respectively (Fig. 8a). ROS levels of *S.* Typhimurium strain was considered as 100%. Data are presented as mean ± S.E.M. (*n* = 4). \*\**p* < 0.01, \*\*\**p <* 0.001

The effect of GSH supplementation on catalase specific activity was determined. Interestingly, we observed 0.70 folds less catalase specific activity in Δ5*msr* mutant cultures supplemented with GSH as compared to the PBS supplemented counterpart (Fig. 8a). Similarly, on native gel, Δ5*msr* mutant cultures supplemented with GSH shows 0.67 folds less KatG activity than the PBS supplemented cultures (Fig. 8b).

**Fig. 8:**
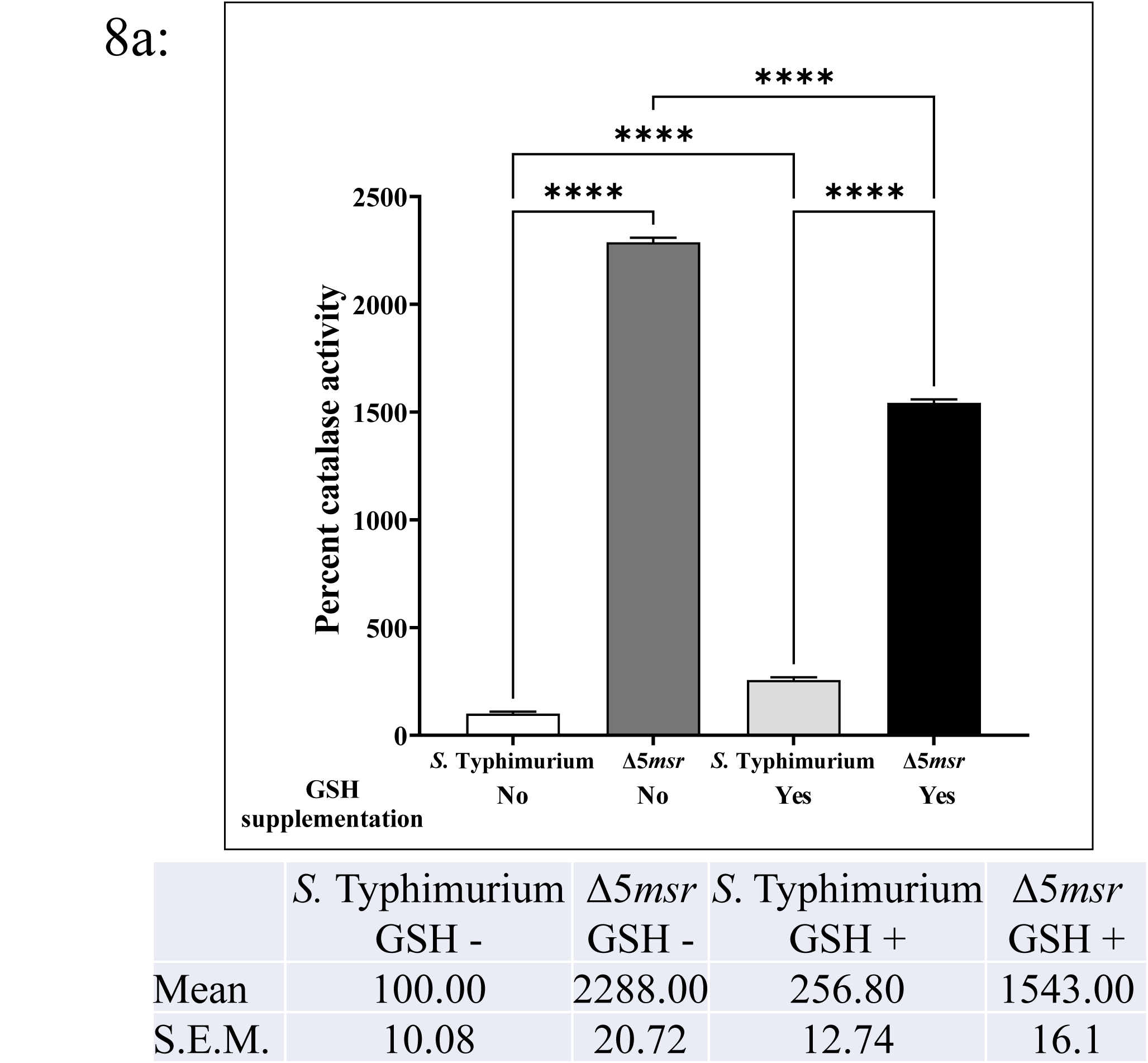
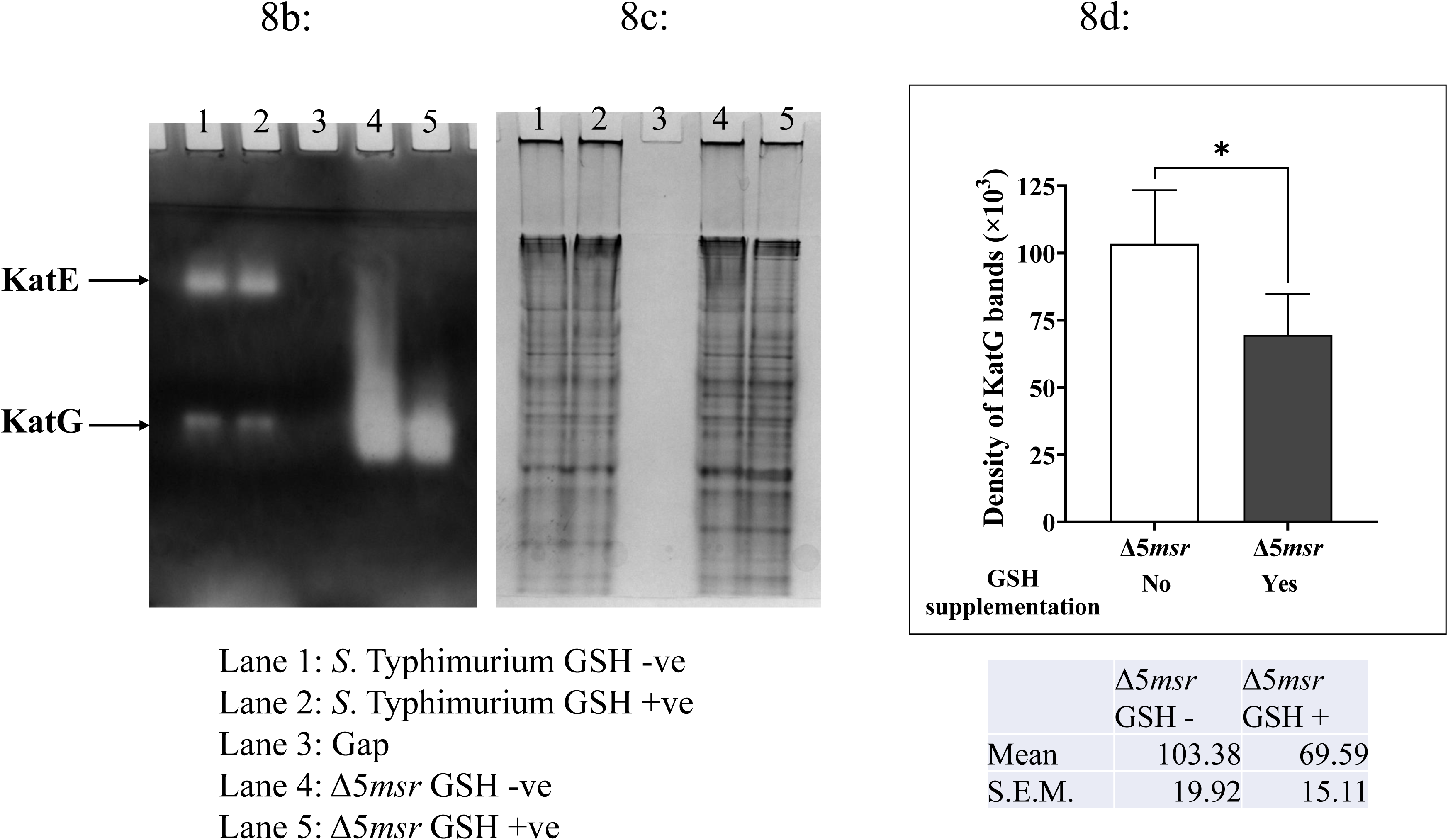
GSH supplemented cultures of Δ5*msr* mutant strain show decreased catalase specific and KatG activities. *S.* Typhimurium and Δ5*msr* mutant strains were cultured with or without supplementation of 10 mM GSH till mid log phase of growth. (A) Catalase specific activity was determined in the cell free lysates as detailed in materials and methods. Activity in *S.* Typhimurium GSH -ve was considered as 100% (Fig. 8a). Data are presented as mean ± S.E.M. (*n* = 6). \*\*\*\**p* < 0.0001 (B) 50 µg cell free lysates of *S.* Typhimurium (GSH –ve & GSH +ve) and Δ5*msr* (GSH –ve & GSH +ve) were resolved on native gel and the catalase activity was determined as described in the material and methods (Fig. 8b). The duplicate native gel stained with CBB (Fig. 8c) served as loading control. The densitometric analysis of bands using ImageJ software is depicted in Fig. 8d. Data are presented as mean ± S.E.M. (*n* = 3). \**p* < 0.05

### GSH supplementation reverses resistance of Δ5*msr* mutant strain against H_2_O_2_

Next, we asked if GSH addition can reverse the resistance of Δ5*msr* mutant strain against H_2_O_2_. Indeed, the GSH supplemented cultures of *S.* Typhimurium and Δ5*msr* mutant strains showed enhanced sensitivity to 25 mM H_2_O_2_ (Fig. 9). The recovered numbers (log_10_ CFU/ml; mean ± S.E.M.) were 3.74 ± 1.90, 2.08 ± 2.08, 8.98 ± 0.10 and 1.10 ± 1.10 for *S.* Typhimurium, *S.* Typhimurium supplemented with GSH, Δ5*msr* mutant strain and Δ5*msr* mutant strain supplemented with GSH, respectively. Furthermore, no viable bacterial cells were recovered when GSH supplemented cultures of Δ5*msr* mutant strain was exposed to 50 mM H_2_O_2_ (Fig. 9).

**Fig. 9:**
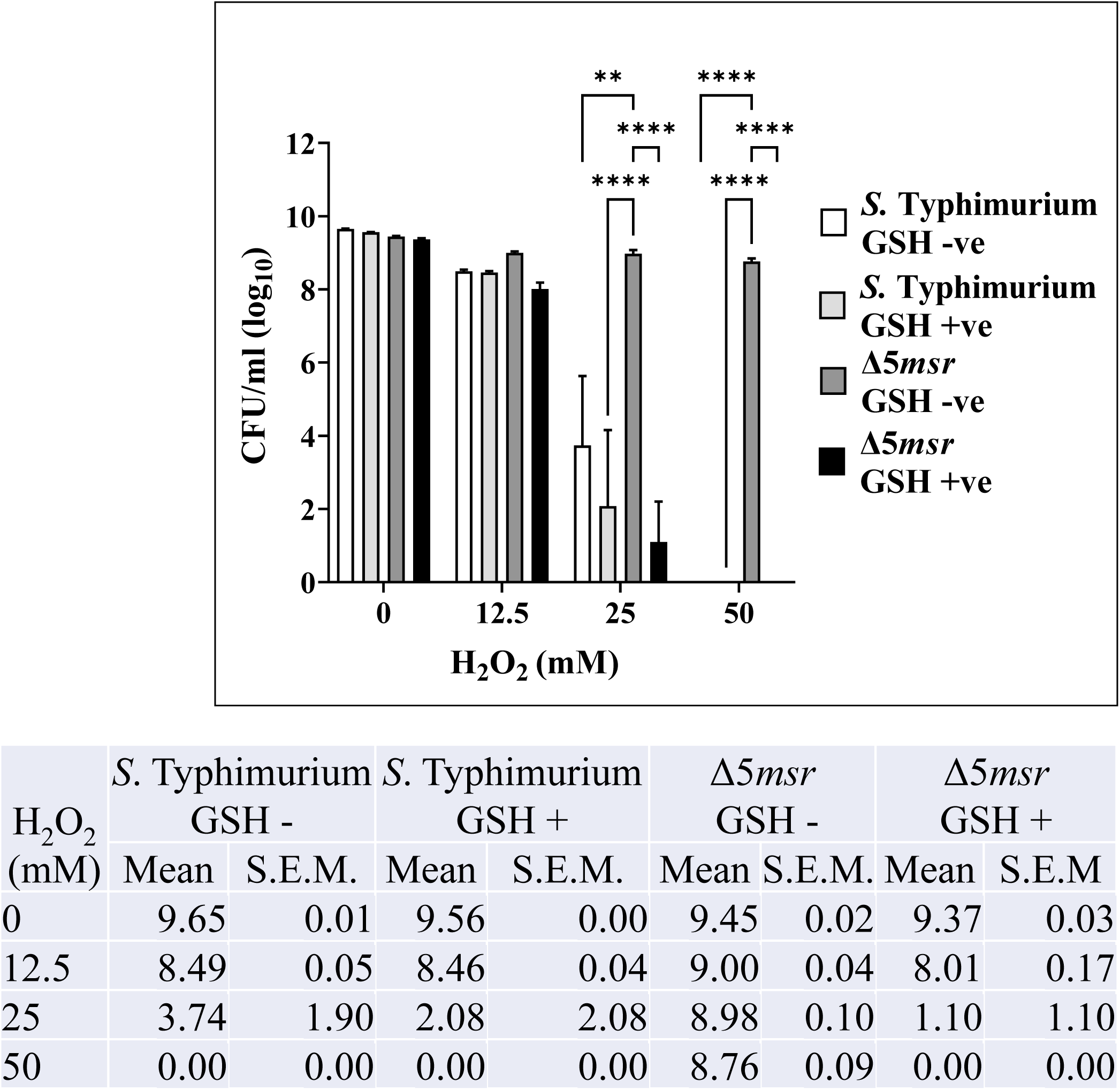
GSH supplementation abrogates resistance of Δ5*msr* mutant strain to H_2_O_2_. GSH supplemented cultures of *S.* Typhimurium and Δ5*msr* mutant strains were exposed to H_2_O_2_ for 2 hours. The cultures were then serially diluted and plated on HE agar plates. The CFUs were counted following overnight incubation of the plates. Data are presented as mean ± S.E.M. (*n* = 3). \*\**p* < 0.01; \*\*\*\**p* < 0.0001

## Discussion

H_2_O_2_ is one of the most important ROS generated in the biological systems. It is not only oxidizes various biomolecules itself but also serves as a substrate for other highly reactive ROS including OH^•^ and HOCl. Therefore, it is not surprising that the bacterial pathogens might have evolved multiple strategies to nullify the effect of H_2_O_2_ (Mishra and Imlay, 2012). Catalases and Msrs are among two important enzymatic systems which have been implicated in H_2_O_2_ resistance.

Contribution of catalases and Msrs in the survival of bacterial pathogens including *S.* Typhimurium under H_2_O_2_ stress has been assessed by mutational analysis. The *katEkatG* double mutant strain of *S.* Typhimurium exhibited increased susceptibility to H_2_O_2_, while the single *katE* or *katG* mutant strains were resistant to H_2_O_2_ (Buchmeier *et al*., 1995). However, several studies reported higher sensitivity of Δ*katE* and Δ*katG* mutant strains of *S.* Typhimurium to H_2_O_2_ (Liao *et al*., 2019; Hahn *et al*., 2021; Kirthika *et al*., 2022) suggesting the important role of catalases in *S.* Typhimurium. The *msrA* and *msrB* double gene deletion (*msrAmsrB* mutant) strain of *Pseudomonas aeruginosa* exhibited hypersensitivity to H_2_O_2_ (Romsang *et al*., 2013). Similarly, *msrA* or *msrB* deletion mutants of *Enterococcus faecalis* (Zhao *et al*., 2010) and *Campylobacter jejuni* (Atack and Kelly, 2008) suffered H_2_O_2_ stress. In *Staphylococcus aureus*, *msrA1* and triple *msrA* (*msrA1, msrA2, msrA3*) mutants have been shown to be hypersensitive to H_2_O_2_ (Singh *et al*., 2015). In *S.* Typhimurium, the *msrA*, *msrC* and *bisC* contribute to the resistance of *S*. Typhimurium against H_2_O_2_ (Denkel *et al*., 2011; Denkel *et al*., 2013).

In current study, first, we generated a pan *msr* (Δ5*msr* mutant) strain (Sahoo et al., 2023). We hypothesized that the lack of all Met-SO repair enzymes in Δ5*msr* mutant strain would result in very high sensitivity of this strain against H_2_O_2_. Contrary to this, to our surprise the Δ5*msr* mutant strain has been more than 2200 fold more resistant to very high concentrations of H_2_O_2_ (25 mM) (Fig. 1). Further, the Δ5*msr* mutant strain depicted very high catalase activity (Fig. 2a).

Among three catalases, KatG and KatE play major role in neutralising H_2_O_2_, while KatN has been shown to have a minor contribution in the survival of *S.* Typhimurium under H_2_O_2_ mediated oxidative stress (Robbe-Saule *et al*., 2001; Kroger *et al*., 2013). To determine the catalase responsible for the very high catalase specific activity in Δ5*msr* mutant strain, we assayed the catalase activity on native gel. Densitometric analysis revealed ∼6 folds higher KatG activity in Δ5*msr* mutant strain (Fig. 2b, 2d). Further, mass spectrometric and transcriptomic analyses revealed enhanced expression of KatG in the Δ5*msr* mutant strain as compared to the *S.* Typhimurium (Supplementary Fig. 1, Fig. 4a). While the expression of *kat*E (Fig. 4b) and *kat*N (Fig. 4c) were found to be downregulated in Δ5*msr* mutant strain.

Transcriptional regulator OxyR regulates several oxidative stress response genes (Chrisman *et al*., 1985; Storz *et al*., 1990). Activation of OxyR depends upon the accumulation of ROS in the cell and subsequent ROS mediated oxidation of Cys_199_ and Cys_208_ residues of OxyR (Zheng and Storz, 1998; Mishra and Imlay, 2012). The oxidized form of OxyR binds to the promoter region of the regulated genes and induces their expression. Glutathione with the help of glutaredoxin1 reduces Cys residues in OxyR and renders the protein inactive (Zheng *and* Storz, 1998; Storz *et al*., 1990; Lee *et al*., 2004). KatG is one of the OxyR regulated gene (Storz *et al*., 1990; Pomposiello and Demple, 2001; Robbe-Saule *et al*., 2001). As Msrs are involved in ROS homeostasis in the cell (Chandra *et al*., 2024) and their absence might accumulate ROS in the cell. Next, we hypothesised that the enhanced expression of KatG might be due to accumulation of ROS in the cell and concomitant oxidation and activation of OxyR. We estimated the ROS levels in Δ5*msr* mutant strain. Indeed, we found more than 5 folds higher ROS accumulation in Δ5*msr* mutant strain (Fig. 5). Further, we argued if upregulation of KatG in Δ5*msr* mutant strain is correlated with ROS levels and OxyR activation, then the neutralisation of ROS might reduce the KatG activity. GSH is known to neutralise ROS and maintain cellular redox balance. We supplemented GSH in the early log growing cultures of *S.* Typhimurium and Δ*5msr* mutant strains and analysed for KatG activity. Indeed, addition of GSH alleviated the ROS levels in Δ5*msr* mutant strain (Fig. 7), along with 0.67 folds reduction in KatG activity (Fig. 8b; 8d). Interestingly, addition of GSH rendered the Δ5*msr* mutant strain sensitive to H_2_O_2_ (Fig. 9).

KatE and KatN are the stationary phase catalases regulated by RpoS (Buchmeier *et al*., 1995; Kroger *et al*., 2013). We have not observed any KatE and KatN activities in the Δ*5msr* mutant strain (Fig. 3a). Previous studies have suggested the inverse relation of oxidizing environment with RpoS induction (Komitopoulou *et al*., 2004). To determine the redox (oxidation) status of Δ*5msr* mutant strain, we compared total thiol levels of *S.* Typhimurium and Δ*5msr* mutant strain. Total thiol levels are being used as an indirect indicator of the general redox (oxidation) status within the cell (Jones and Go, 2010). The Δ*5msr* mutant strain showed significantly lower total thiols than *S*. Typhimurium (Fig. 6). The higher ROS and lower total thiols levels suggest high oxidative environment in absence of *msr* which might be preventing activation of RpoS, and thus preventing activation of these two RpoS regulated catalases.

Our findings suggest the ROS dependent upregulation of KatG is associated with very high resistance of the Δ*5msr* mutant strain against H_2_O_2_. *msr* gene deletion strains of few bacterial pathogens showed resistance against H_2_O_2_ stress. For example, *msrB* and quadruple *msr* (*msrA1, msrA2, msrA3, msrB*) deletion strains of *Staphylococcus aureus* showed resistance to H_2_O_2_ (Singh *et al*., 2015). Similarly, *msrB* and *msrAmsrB* double gene deletion strains of *Mycobacterium smegmatis* showed resistance to H_2_O_2_ (Dhandayuthapani *et al*., 2009). These studies have not elucidated the mechanism of H_2_O_2_ resistance in *msr* mutant strain. However, it would be very interesting to know if the mechanism of H_2_O_2_ resistance in these cases is catalase dependent.

Interestingly, mice lacking the entire Msr repair system (quadruple *msr* knockout mice) displayed resistance to oxidative stress (induced by ischemic reperfusion and paraquat exposure), suggesting that the absence of Msrs trigger adaptive resistance mechanisms (Lai *et al*., 2019). Similarly, murine embryonic fibroblasts derived from quadruple *msr* knockout mice showed similar sensitivity to H_2_O_2_ and paraquat as compared to wild-type cells (Lai *et al*., 2019). Additionally, Hepatocyte cell line ALM21 lacking MsrB with heterozygous MsrA

(Msrb^3KO^ -Msra+/-) and wild type cells exhibited similar susceptibility to ischemia-reperfusion and paraquat (Trujillo-Hernandez and Levine, 2023). These findings clearly highlight the induction of adaptive resistance mechanism(s) involved in cellular survival in the absence of Msrs. However, these studies have not elucidated/ identified the factor involved in such resistance.

The importance/co-operative roles of KatG and Msrs in managing oxidative stress and maintaining bacterial survival are further underscored by our inability to delete *katG* from the Δ5*msr* mutant strain background. However, as a control we were able to delete *katG* from *S.* Typhimurium (wild type) strain. The deletion of *katG* was confirmed by a PCR (Supplementary Fig. S2). Additionally, catalase activity on native gels did not show any KatG activity in the *katG* deletion mutant strain (Supplementary Fig. S3). This indicates that KatG is essential for bacterial growth and survival in the absence of *msr*s.

Interestingly, KatG is shown to be involved in antibiotic susceptibility. In *Mycobacterium tuberculosis*, KatG is shown to activate isoniazid (INH) as Δ*katG* mutant strain showed INH resistance (Zhang *et al*., 1992; Hazbon *et al*., 2006). While KatG reduces kanamycin susceptibility in *E. coli* (Loewen *et al*., 2018). Therefore, modulation in expression of KatG might be targeted for development of antibacterial therapies.

In conclusion, current study reveals the critical compensatory role of KatG in managing oxidative stress in *S.* Typhimurium when the Msr repair system is absent. Contrary to our initial hypothesis that the lack of *msr* would increase sensitivity to peroxide, the Δ5*msr* mutant strain exhibited remarkable resistance to high concentrations of H_2_O_2_, primarily due to a significant upregulation of KatG activity. This upregulation is ROS dependent, likely mediated by the oxidative activation of OxyR, and suggests a complex adaptive response where one ROS neutralising mechanism compensates for the loss of another. The absence of Msr created an oxidizing intracellular environment, reflected in increased ROS accumulation and lower thiols levels, which further influenced the inactivation of RpoS, leading to reduced activities of KatE and KatN. This study underscores the importance of the complex interplay between different bacterial defence systems in response to oxidative stress.

## Experimental procedures

### Bacterial strains and culture conditions

The *Salmonella enterica* subspecies *enterica* serovar Typhimurium strain E-5591 (*S.* Typhimurium) was obtained from the National *Salmonella* Centre, ICAR-IVRI, Izatnagar, India. Δ*5msr* mutant strain was cured as described elsewhere (Sahoo *et al*., 2023). *S.* Typhimurium was cultured on Hektoen enteric (HE) agar or in Luria Bertani (LB) broth. KatG deletion strain was constructed as per the protocol described elsewhere (Sarkhel *et al*., 2017; Pesingi *et al*., 2017; Datsenko and Wanner, 2000). The primers are detailed in Supplementary Table S2.

### H_2_O_2_ sensitivity assays

The overnight grown cultures of *S.* Typhimurium and Δ5*msr* mutant strains were diluted in LB broth at a ratio of 1:100 and grown till the mid log phase at 37 °C. The mid log grown cultures were exposed to various concentrations of H_2_O_2_ at 37 °C for 2 hours. The cultures were then serially diluted and plated on HE agar plates. The CFUs were counted following overnight incubation of the plates. In few experiments, the cultures were supplemented with reduced GSH (10 mM) at 0.3 OD (early log phase).

### Preparation of cell-free lysates

Early (OD_600_ 0.7), mid (OD_600_ 0.8 – 1.0), late log (OD_600_ 1.3-1.4), and stationary phase (OD_600_ 2.5) grown cultures were pelleted at 4500 x *g* for 15 min at 4 °C. The pellets were washed twice with ice cold PBS and lysed by 15 cycles of sonication. The lysates were centrifuged at 14000 x *g* for 30 min at 4 °C. The supernatants were collected and stored at -80 °C in aliquots. Total proteins in such cell-free lysates were estimated by Pierce BCA Protein Assay Kit (Thermo Scientific).

### Estimation of catalase specific activity

Catalase specific activity in the cell free lysates was determined spectrophotometrically by measuring the dissipation of H_2_O_2_ at 240 nm (Aebi, 1984; Mahawar *et al*., 2011). The catalase specific activity was calculated using Beer Lambert’s law, taking 43.6 mol^-1^ cm^-1^ as the molar absorption coefficient of H_2_O_2_ (Noble and Gibson, 1970; McLean *et al*., 2010).

### Analysis of catalase activity on native gel

Catalase activity on native gel was analysed as described elsewhere (Weydert and Cullen, 2010). Briefly, the 50 µg of cell-free lysates were resolved on 10% native polyacrylamide gels. After 2 washes with distilled water, the gels were incubated in 2 mM of H_2_O_2_ and subsequently stained with 2% potassium ferricyanide and 2% ferric chloride. The staining solution was removed as soon as achromatic bands appeared. The gels were washed, scanned, and analysed by ImageJ.

### Mass spectrometry analysis

Mid log grown cultures of *S.* Typhimurium and Δ5*msr* mutant strains were pelleted and boiled in 2X SDS gel-loading dye containing β-mercaptoethanol. Samples were resolved in 10% SDS gel and stained with CBB. The upregulated protein band was identified by peptide mass fingerprinting followed by MASCOT analysis by outsourcing to Biotech Desk Pvt. Ltd, Hyderabad, India.

### Complementary DNA (cDNA) synthesis and real-time quantitative polymerase chain reaction (RT-qPCR)

0.4, 0,7 and 1.0 OD_600_ grown cultures of *S.* Typhimurium and Δ5*msr* mutant strains were pelleted. Total RNA was extracted using RNAiso Plus (Takara Bio, Shiga, Japan) and precipitated with isopropanol following a standard protocol (Chomczynski and Sacchi, 1987). Subsequently, cDNA synthesis was performed in a 20-μl reaction volume using the Maxima H minus First Strand cDNA Synthesis Kit (Thermo Scientific) as per the manufacturer’s instructions. RT-qPCR was conducted using 2X Maxima SYBR Green/ROX qPCR master mix (Thermo Scientific) in the Applied Biosystems™ StepOne real-time PCR system. The real time PCR data were analysed by 2^-ΔΔCT^ method (Livak and Schmittgen, 2001) where, ΔΔCT = (CT_Target gene_ –CT_gmk_)_Δ5*msr*_ - (CT_Target gene_ – CT_gmk_)*_S._* _Typhimurium_. The relative fold-change in mRNA expression levels of *kat*G, *kat*E, and *kat*N was determined using specific primers listed in Supplementary Table S1.

### Estimation of total thiols

Total thiol levels were estimated using 5,5’-dithio-bis-(2-nitrobenzoic acid) (DTNB) as previously described (Ellman, 1959) with minor modifications. Briefly, PBS, cell-free lysates and 2% SDS were mixed thoroughly in a cuvette. The reaction was initiated by addition of 100 µM DTNB. The increase in absorbance with the production of 5-thio-2- nitrobenzoic acid (TNB) was recorded at 412 nm. The molar extinction coefficient of TNB at 412 nm is 13.8 × 10^3^ M^-1^ cm^-1^ (Eyer *et al*., 2003).

### Quantitation of intracellular ROS levels

Intracellular ROS levels were estimated by 2’,7’-dichlorodihydrofluorescein diacetate (DCFDA) as described elsewhere (Chandra *et al*., 2024). Briefly, vigorously growing cultures of *S.* Typhimurium and Δ5*msr* mutant strains were pelleted, washed, suspended in PBS and incubated with 10 μM DCFDA for 30 min in dark. The cells were then washed two times with PBS and suspended in 1 ml of PBS. 200 μl of these suspensions were added in black 96 well plates (Thermo Fisher Immuno Standard Modules Black). The plate was read at excitation of 485 nm and emission of 535 nm using SpectraMax M5 fluorometer (Molecular devices). The fluorescence without addition of DCFDA were subtracted. The fluorescence in DCFDA exposed *S.* Typhimurium cultures were considered 100%. Simultaneously, the DCFDA exposed cultures were spotted on glass slides and observed under a fluorescent microscope (Nikon ECLIPSE T*i –* S) using a green fluorescent filter.

### Statistical analysis

Data were analysed using GraphPad Prism version 9.3.1 (Trial version). Comparisons between multiple groups were done using one-way analysis of variance (ANOVA) or two-way ANOVA followed by Tukey’s post-hoc test or paired t-test. *p* values <0.05 were considered as significant.

## Authors’ Contributions

L.L., S.U., R.S., T.K.S.C., and M.M. designed the experiments. L.L., S.U., R.S., and T.K.S.C. performed the experiments and analyzed the data. L.L. and M.M. wrote and edited the manuscript. All authors read and approved the final manuscript.

## Additional Information

The authors declare no competing interests.

## Acknowledgements

We acknowledge the financial support provided by Department of Biotechnology (DBT), Govt. of India, Grant number BT/PR46085/AAQ/1/894/2022. We thank Dr. Robert J. Maier, Department of Microbiology, University of Georgia, Athens, GA, USA for his kind gift of the plasmids pKD3, pKD46, and pCP20. Our gratitude is extended to the Director of ICAR-Indian Veterinary Research Institute, Izatnagar, Bareilly, India for the provision of necessary facilities.

## Data availability statement

The data generated and analyzed in the current study are available with the corresponding author.

---

**Fig. 13:**
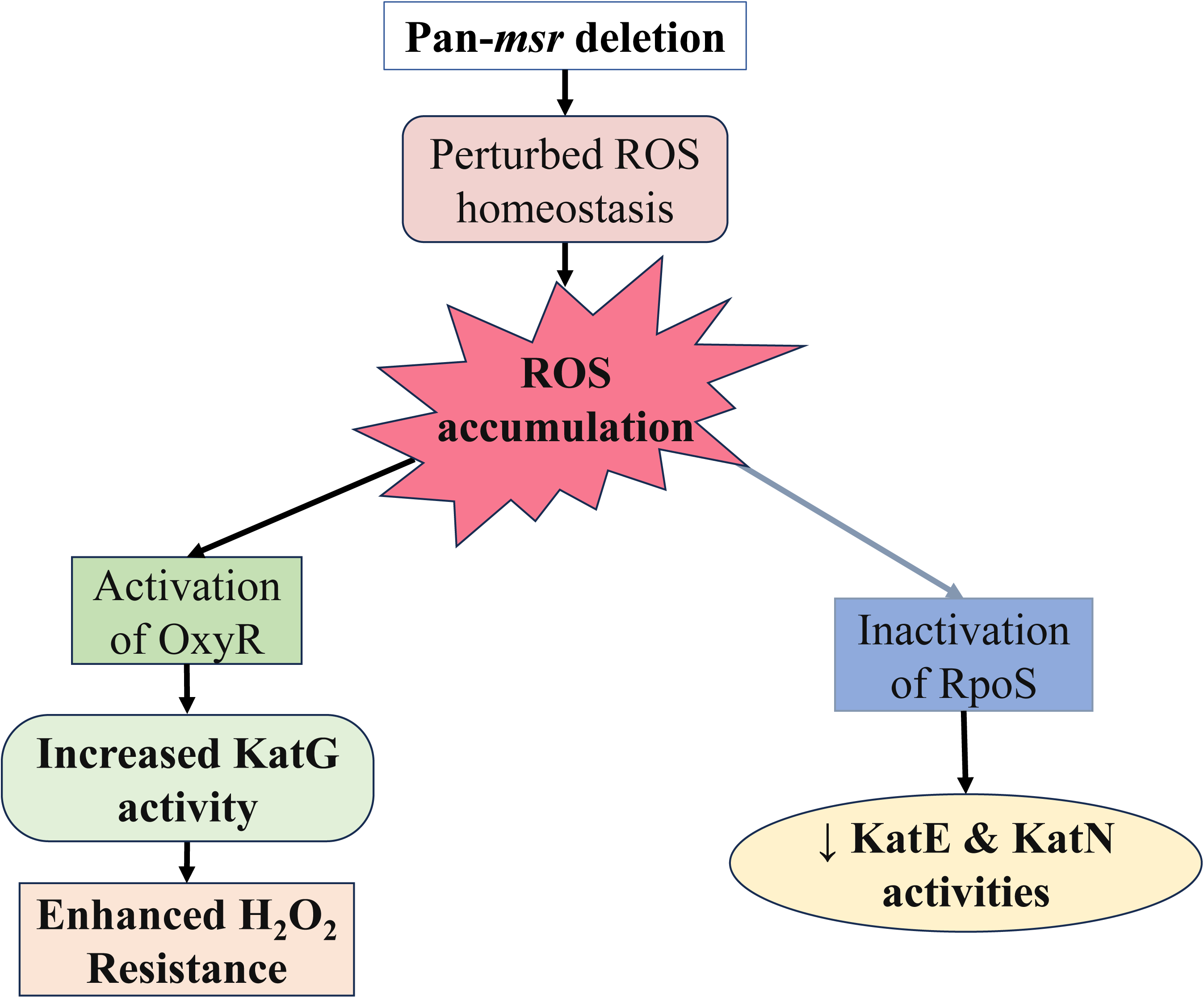
Proposed model. The deletion of *msrs* leads to perturbed ROS homeostasis and subsequent accumulation of ROS in the cell. The ROS might oxidise and activate OxyR, resulting in enhanced KatG activity. The increased KatG activity enhances the survival of Δ5*msr* mutant strain against H_2_O_2_. However, GSH by neutralizing ROS, decreases KatG activity and subsequently abrogate H_2_O_2_ resistance. Further, high ROS levels create an oxidizing environment, reduces levels of thiols that likely inhibits RpoS, decreasing the KatE and KatN activities in mutant strain.

**Supplementary Fig.S1:**
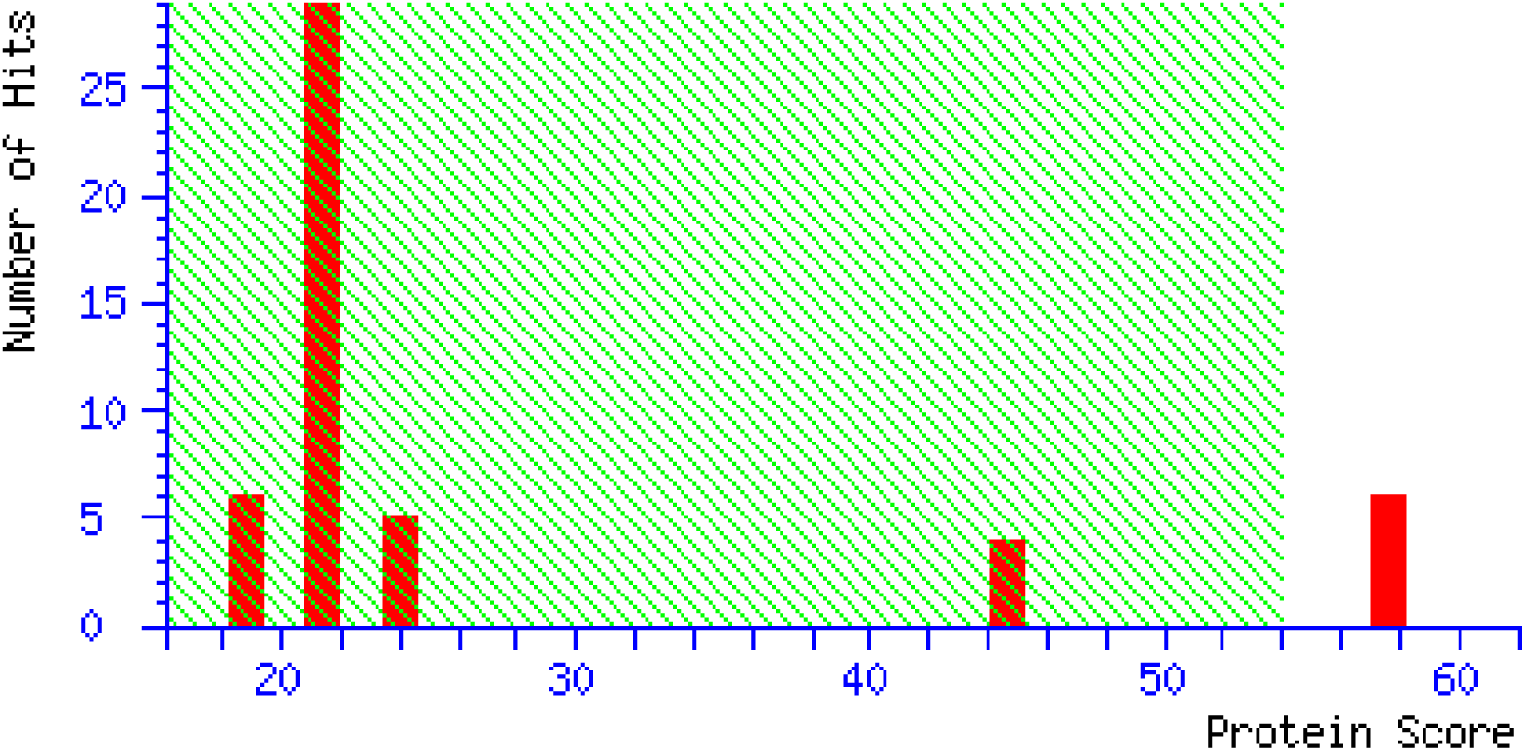
The upregulated protein band observed in the SDS gel was identified by peptide mass fingerprinting followed by MASCOT analysis. The protein has a score of 58.

**Supplementary Fig. S2:**
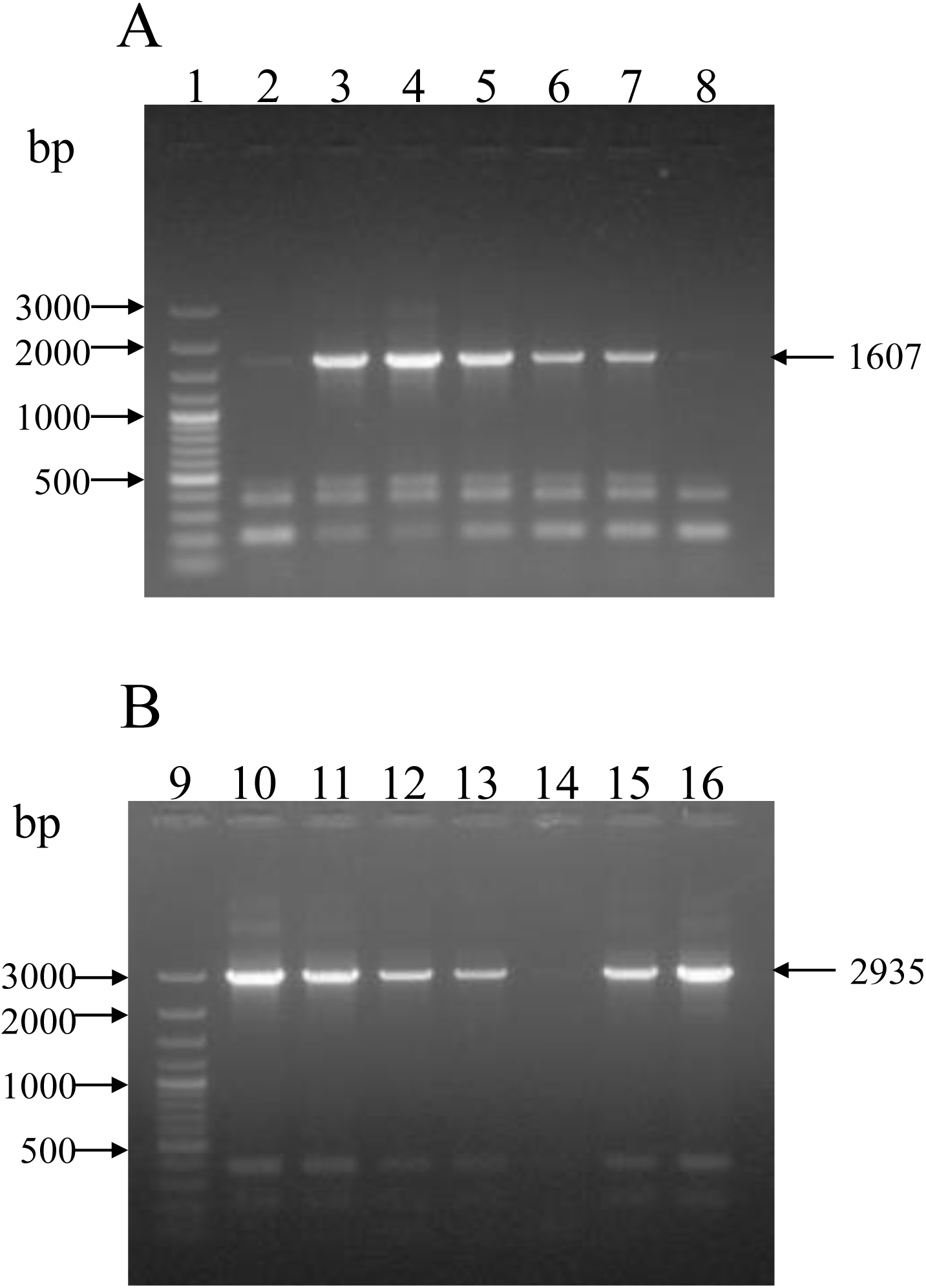
The *katG* was deleted either in *S.* Typhimurium (Fig. A) or in Δ5*msr* mutant strain (Fig. B) background using the method described elsewhere (Datsenko and Wanner, 2000). Lane 1 and 9 are 100 bp plus DNA ladder. PCR product obtained using flanking primers and Δ*katG* mutant in *S.* Typhimurium background (lane 2 to 8) or Δ5*msr* mutant background (lane 10 to 16). Expected size of amplicon if *katG* deleted = 1607 bp and 2935 bp if *katG* is not deleted.

**Supplementary Fig. S3:**
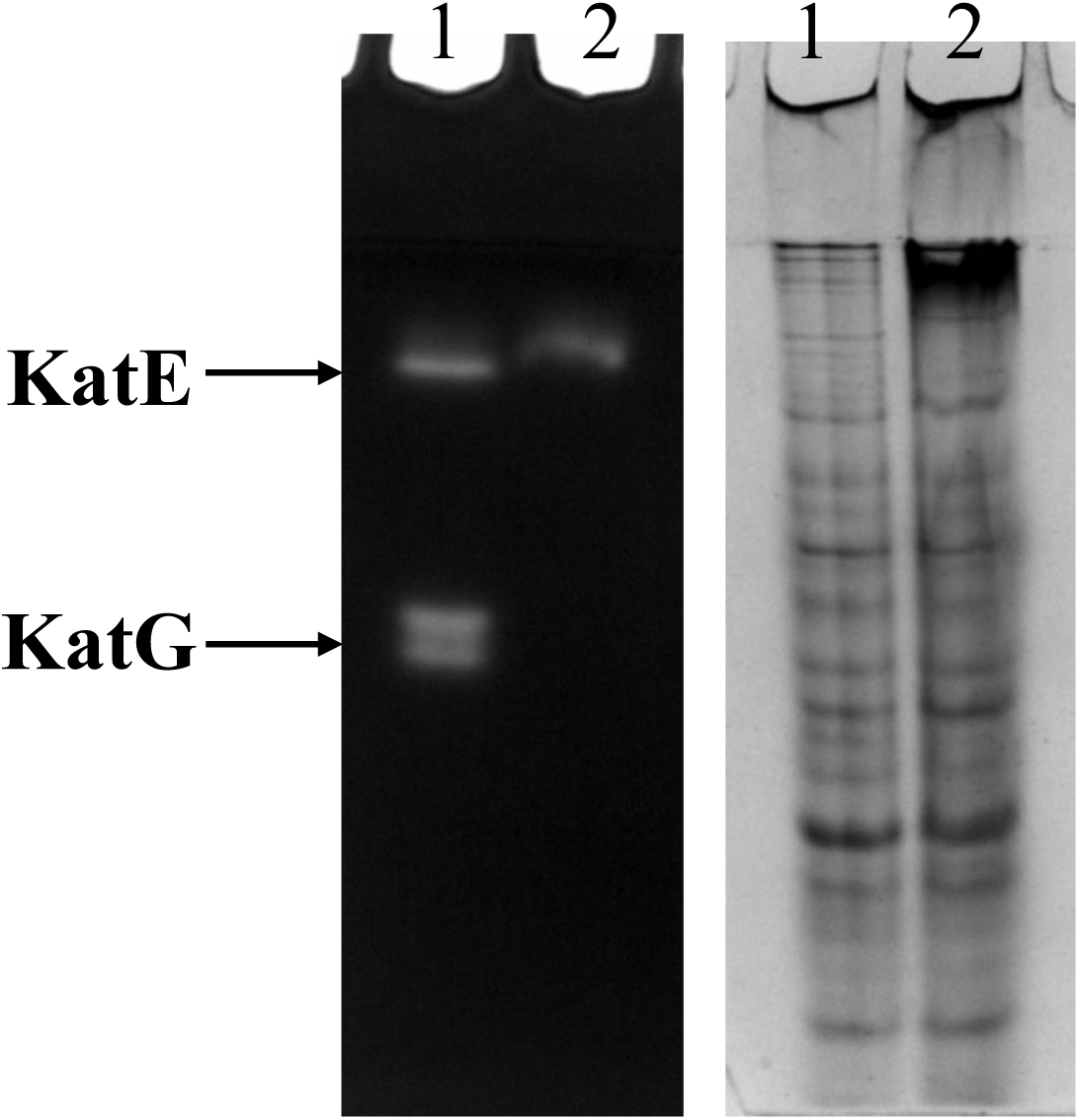
50 µg cell free lysates of *S.* Typhimurium and Δ*katG* mutant strains were resolved on native gel and the catalase activity was determined as described in the material and methods. The duplicate native gel stained with CBB served as a loading control.

## References

1. CDC Yellow Book 2024: Health Information for International Travel (New York, 2023; online edn, Oxford Academic, 23 Mar. 2023).

2. Gilchrist, J. J., & MacLennan, C. A. (2019). Invasive Nontyphoidal Salmonella Disease in Africa. EcoSal Plus, 8(2), 10.1128/ecosalplus.ESP-0007-2018.

3. Balasubramanian R., Im J., Lee J. S., Jeon H. J., Mogeni O. D., Kim J. H., et al. (2019). The global burden and epidemiology of invasive non-typhoidal *Salmonella* infections. Hum. Vaccin. Immunother. 15, 1421–1426. doi: 10.1080/21645515.2018.1504717

4. Imlay, J. A. (2013). The molecular mechanisms and physiological consequences of oxidative stress: Lessons from a model bacterium. Nature Reviews Microbiology, 11(7), 443–454. 10.1038/nrmicro3032

5. Seixas, A. F., Quendera, A. P., Sousa, J. P., Silva, A. F. Q., Arraiano, C. M., & Andrade, J. M. (2022). Bacterial Response to Oxidative Stress and RNA Oxidation. Frontiers in Genetics, 12(January), 1–12. 10.3389/fgene.2021.821535

6. Kehrer J. P. (2000). The Haber-Weiss reaction and mechanisms of toxicity. Toxicology, 149(1), 43–50. 10.1016/s0300-483x(00)00231-6

7. Fasnacht, M., & Polacek, N. (2021). Oxidative Stress in Bacteria and the Central Dogma of Molecular Biology. Frontiers in Molecular Biosciences, 8(May), 1–13. 10.3389/fmolb.2021.671037

8. Levine, R. L., Mosoni, L., Berlett, B. S., & Stadtman, E. R. (1996). Methionine residues as endogenous antioxidants in proteins. Proceedings of the National Academy of Sciences, 93(26), 15036–15040.

9. Fang, F. C., Degroote, M. A., Foster, J. W., Bäumler, A. J., Ochsner, U., Testerman, T., Bearson, S., Giárd, J. C., Xu, Y., Campbell, G., & Laessig, T. (1999). Virulent Salmonella typhimurium has two periplasmic Cu, Zn-superoxide dismutases. Proceedings of the National Academy of Sciences of the United States of America, 96(13), 7502–7507. 10.1073/pnas.96.13.7502

10. Robbe-Saule, V., Coynault, C., Ibanez-Ruiz, M., Hermant, D., & Norel, F. (2001). Identification of a non-haem catalase in Salmonella and its regulation by RpoS (σs). Molecular Microbiology, 39(6), 1533–1545. 10.1046/j.1365-2958.2001.02340.x

11. Hébrard, M., Viala, J. P. M., Méresse, S., Barras, F., & Aussel, L. (2009). Redundant hydrogen peroxide scavengers contribute to Salmonella virulence and oxidative stress resistance. Journal of Bacteriology, 191(14), 4605–4614. 10.1128/JB.00144-09

12. Kröger, C., Colgan, A., Srikumar, S., Händler, K., Sivasankaran, S. K., Hammarlöf, D. L., Canals, R., Grissom, J. E., Conway, T., Hokamp, K., & Hinton, J. C. D. (2013). An infection-relevant transcriptomic compendium for salmonella enterica serovar typhimurium. Cell Host and Microbe, 14(6), 683–695. 10.1016/j.chom.2013.11.010

13. Hillar, A., Peters, B., Pauls, R., Loboda, A., Zhang, H., Mauk, A. G., & Loewen, P. C. (2000). Modulation of the Activities of Catalase - Peroxidase HPI of Escherichia coli by. Biochemistry, 39(19), 5868–5875.

14. Mishra, S., & Imlay, J. (2012). Why do bacteria use so many enzymes to scavenge hydrogen peroxide?. Archives of biochemistry and biophysics, 525(2), 145–160. 10.1016/j.abb.2012.04.014

15. Hahn, M. M., González, J. F., & Gunn, J. S. (2021). Salmonella Biofilms Tolerate Hydrogen Peroxide by a Combination of Extracellular Polymeric Substance Barrier Function and Catalase Enzymes. Frontiers in Cellular and Infection Microbiology, 11(May), 1–15. 10.3389/fcimb.2021.683081

16. Kirthika, P., Jawalagatti, V., Senevirathne, A., & Lee, J. H. (2022). Coordinated interaction between Lon protease and catalase-peroxidase regulates virulence and oxidative stress management during Salmonellosis. Gut Microbes, 14(1). 10.1080/19490976.2022.2064705

17. McLean, S., Bowman, L. A. H., & Poole, R. K. (2010). KatG from Salmonella Typhimurium is a peroxynitritase. FEBS Letters, 584(8), 1628–1632. 10.1016/j.febslet.2010.03.029

18. Luan, G., Hong, Y., Drlica, K., & Zhao, X. (2018). Suppression of Reactive Oxygen Species Accumulation Accounts for Paradoxical Bacterial Survival at High Quinolone Concentration. Antimicrobial agents and chemotherapy, 62(3), e01622–17. 10.1128/AAC.01622-17

19. Mahawar, M., Tran, V. L., Sharp, J. S., & Maier, R. J. (2011). Synergistic roles of Helicobacter pylori methionine sulfoxide reductase and GroEl in repairing oxidant-damaged catalase. Journal of Biological Chemistry, 286(21), 19159–19169. 10.1074/jbc.M111.223677

20. Benoit, S. L., Bayyareddy, K., Mahawar, M., Sharp, J. S., & Maier, R. J. (2013). Alkyl hydroperoxide reductase repair by Helicobacter pylori methionine sulfoxide reductase. Journal of Bacteriology, 195(23), 5396–5401. 10.1128/JB.01001-13

21. Kuhns, L. G., Mahawar, M., Sharp, J. S., Benoit, S., & Maier, R. J. (2013). Role of Helicobacter pylori methionine sulfoxide reductase in urease maturation. Biochemical Journal, 450(1), 141–148. 10.1042/BJ20121434

22. Gennaris, A., Ezraty, B., Henry, C., Agrebi, R., Vergnes, A., Oheix, E., Bos, J., Leverrier, P., Espinosa, L., Szewczyk, J., Vertommen, D., Iranzo, O., Collet, J. F., & Barras, F. (2015). Repairing oxidized proteins in the bacterial envelope using respiratory chain electrons. Nature, 528(7582), 409–412. 10.1038/nature15764

23. Sarkhel, R., Rajan, P., Gupta, A. K., Kumawat, M., Agarwal, P., Shome, A., Puii, L., & Mahawar, M. (2017). Methionine sulfoxide reductase A of Salmonella Typhimurium interacts with several proteins and abets in its colonization in the chicken. Biochimica et Biophysica Acta - General Subjects, 1861(12), 3238–3245. 10.1016/j.bbagen.2017.09.014

24. Andrieu, C., Loiseau, L., Vergnes, A., Gagnot, S., Barré, R., Aussel, L., … & Ezraty, B. (2023). Salmonella Typhimurium uses the Cpx stress response to detect N-chlorotaurine and promote the repair of oxidized proteins. Proceedings of the National Academy of Sciences, 120(14), e2215997120.

25. Denkel, L. A., Horst, S. A., Rouf, S. F., Kitowski, V., Böhm, O. M., Rhen, M., Jäger, T., & Bange, F. C. (2011). Methionine sulfoxide reductases are essential for virulence of Salmonella typhimurium. PLoS ONE, 6(11). 10.1371/journal.pone.0026974

26. Denkel, L. A., Rhen, M., & Bange, F. C. (2013). Biotin sulfoxide reductase contributes to oxidative stress tolerance and virulence in Salmonella enterica serovar Typhimurium. Microbiology (United Kingdom), 159(PART7), 1447–1458. 10.1099/mic.0.067256-0

27. Trivedi, R. N., Agarwal, P., Kumawat, M., Pesingi, P. K., Gupta, V. K., Goswami, T. K., & Mahawar, M. (2015). Methionine sulfoxide reductase A (MsrA) contributes to Salmonella Typhimurium survival against oxidative attack of neutrophils. Immunobiology, 220(12), 1322–1327. 10.1016/j.imbio.2015.07.011

28. Nair, S. S., Chauhan, T. K. S., Kumawat, M., Sarkhel, R., Apoorva, S., Shome, A., Athira, V., Kumar, B., Abhishek, & Mahawar, M. (2021). Deletion of both methionine sulfoxide reductase A and methionine sulfoxide reductase C genes renders Salmonella Typhimurium highly susceptible to hypochlorite stress and poultry macrophages. Molecular Biology Reports, 48(4), 3195–3203. 10.1007/s11033-021-06381-2

29. Andrieu, C., Vergnes, A., Loiseau, L., Aussel, L., & Ezraty, B. (2020). Characterisation of the periplasmic methionine sulfoxide reductase (MsrP) from Salmonella Typhimurium. Free Radical Biology and Medicine, 160(August), 506–512. 10.1016/j.freeradbiomed.2020.06.031

30. Shome, A., Kumawat, M., Pesingi, P. K., Bhure, S. K., & Mahawar, M. (2020). Isolation and identification of periplasmic proteins in Salmonella Typhimurium. Int. J. Curr. Microbiol. Appl. Sci, 9, 1923–1936.

31. Chandra, H. B., Shome, A., Sahoo, R., Apoorva, S., Bhure, S. K., & Mahawar, M. (2023). Periplasmic methionine sulfoxide reductase (MsrP)—a secondary factor in stress survival and virulence of *Salmonella* Typhimurium. FEMS Microbiology Letters, 370, fnad063. 10.1093/femsle/fnad063

32. Dixit, S. K., Hota, D. P., Rajan, P., Mishra, P. K. K., Goswami, T. K., & Mahawar, M. (2017). *Salmonella* Typhimurium methionine sulfoxide reductase A (MsrA) prefers TrxA in repairing methionine sulfoxide. Preparative Biochemistry and Biotechnology, 47(2), 137–142. 10.1080/10826068.2016.1185733

33. 33. Ezraty, B., Gennaris, A., Barras, F., & Collet, J. (2017). Oxidative stress, protein damage and repair in bacteria. Nature Publishing Group. 10.1038/nrmicro.2017.26

34. Ezraty, B., Grimaud, R., El Hassouni, M., Moinier, D., & Barras, F. (2004). Methionine sulfoxide reductases protect Ffh from oxidative damages in Escherichia coli. The EMBO journal, 23(8), 1868–1877. 10.1038/sj.emboj.7600172

35. Khor, H. K., Fisher, M. T., & Schöneich, C. (2004). Potential role of methionine sulfoxide in the inactivation of the chaperone GroEL by hypochlorous acid (HOCl) and peroxynitrite (ONOO–). Journal of Biological Chemistry, 279(19), 19486–19493.

36. Nasreen, M., Nair, R. P., McEwan, A. G., & Kappler, U. (2022). The Peptide Methionine Sulfoxide Reductase (MsrAB) of *Haemophilus influenzae* Repairs Oxidatively Damaged Outer Membrane and Periplasmic Proteins Involved in Nutrient Acquisition and Virulence. *Antioxidants (Basel*, Switzerland), 11(8), 1557. 10.3390/antiox11081557

37. Levine, R. L., Berlett, B. S., Moskovitz, J., Mosoni, L., & Stadtman, E. R. (1999). Methionine residues may protect proteins from critical oxidative damage. Mechanisms of Ageing and Development, 107(3), 323–332. 10.1016/S0047-6374(98)00152-3

38. Abulimiti, A., Qiu, X., Chen, J., Liu, Y., & Chang, Z. (2003). Reversible methionine sulfoxidation of Mycobacterium tuberculosis small heat shock protein Hsp16.3 and its possible role in scavenging oxidants. Biochemical and biophysical research communications, 305(1), 87–93. 10.1016/s0006-291x(03)00685-5

39. Luo, S., & Levine, R. L. (2009). Methionine in proteins defends against oxidative stress. The FASEB Journal, 23(2), 464.

40. Benoit, S. L., & Maier, R. J. (2016). Helicobacter catalase devoid of catalytic activity protects the Bacterium against oxidative stress. Journal of Biological Chemistry, 291(45), 23366–23373. 10.1074/jbc.M116.747881

41. Schmalstig, A. A., Benoit, S. L., Misra, S. K., Sharp, J. S., & Maier, R. J. (2018). Noncatalytic Antioxidant Role for Helicobacter pylori Urease. Journal of bacteriology, 200(17), e00124–18. 10.1128/JB.00124-18

42. Stadtman, E. R., Moskovitz, J., & Levine, R. L. (2003). Oxidation of methionine residues of proteins: biological consequences. Antioxidants and Redox Signaling, 5(5), 577–582.

43. Sahoo, R., Chauhan, T. K. S., Lalhmangaihzuali, L., Sinha, E., Qureshi, S., & Mahawar, M. (2023). Pan *msr* gene deleted strain of *Salmonella* Typhimurium suffers oxidative stress, depicts macromolecular damage and attenuated virulence. Scientific Reports, 13(1), 1–14. 10.1038/s41598-023-48734-w

44. Mastroeni, P., Vazquez-Torres, A., Fang, F. C., Xu, Y., Khan, S., Hormaeche, C. E., & Dougan, G. (2000). Antimicrobial actions of the NADPH phagocyte oxidase and inducible nitric oxide synthase in experimental salmonellosis. II. Effects on microbial proliferation and host survival in vivo. The Journal of experimental medicine, 192(2), 237–248.

45. St. John, G., Brot, N., Ruan, J., Erdjument-Bromage, H., Tempst, P., Weissbach, H., & Nathan, C. (2001). Peptide methionine sulfoxide reductase from *Escherichia coli* and *Mycobacterium tuberculosis* protects bacteria against oxidative damage from reactive nitrogen intermediates. Proceedings of the National Academy of Sciences of the United States of America, 98(17), 9901–9906. 10.1073/pnas.161295398

46. Alamuri, P., & Maier, R. J. (2006). Methionine sulfoxide reductase in *Helicobacter pylori*: Interaction with methionine-rich proteins and stress-induced expression. Journal of Bacteriology, 188(16), 5839–5850. 10.1128/JB.00430-06

47. Vattanaviboon, P., Seeanukun, C., Whangsuk, W., Utamapongchai, S., & Mongkolsuk, S. (2005). Important role for methionine sulfoxide reductase in the oxidative stress response of *Xanthomonas campestris pv. phaseoli*. Journal of Bacteriology, 187(16), 5831–5836. 10.1128/JB.187.16.5831-5836.2005

48. Atack, J. M., & Kelly, D. J. (2009). Oxidative stress in *Campylobacter jejuni*: responses, resistance and regulation. Future microbiology, 4(6), 677–690.

49. Lei, Y., Zhang, Y., Guenther, B. D., Kreth, J., & Herzberg, M. C. (2011). Mechanism of adhesion maintenance by methionine sulphoxide reductase in *Streptococcus gordonii*. Molecular Microbiology, 80(3), 726–738. 10.1111/j.1365-2958.2011.07603.x

50. Jalal, N., & Lee, S. F. (2020). The MsrAB reducing pathway of *Streptococcus gordonii* is needed for oxidative stress tolerance, biofilm formation, and oral colonization in mice. PLoS ONE, 15(2), 1–19. 10.1371/journal.pone.0229375

51. Weydert, C. J., & Cullen, J. J. (2010). Measurement of superoxide dismutase, catalase and glutathione peroxidase in cultured cells and tissue. Nature protocols, 5(1), 51–66. 10.1038/nprot.2009.197

52. Chandra, H. B., Lalhmangaihzuali, L., Shome, A., Sahoo, R., Irungbam, K., & Mahawar, M. (2024). Comparative analysis reveals the trivial role of MsrP in defending oxidative stress and virulence of Salmonella Typhimurium in mice. Free radical biology & medicine, 213, 322–326. 10.1016/j.freeradbiomed.2024.01.020

53. Kwon, D. H., Cha, H. J., Lee, H., Hong, S. H., Park, C., Park, S. H., Kim, G. Y., Kim, S., Kim, H. S., Hwang, H. J., & Choi, Y. H. (2019). Protective effect of glutathione against oxidative stress-induced cytotoxicity in RAW 264.7 macrophages through activating the nuclear factor erythroid 2-related factor-2/heme oxygenase-1 pathway. Antioxidants, 8(4). 10.3390/antiox8040082

54. Buchmeier, N. A., Libby, S. J., Xu, Y., Loewen, P. C., Switala, J., Guiney, D. G., & Fang, F. C. (1995). DNA repair is more important than catalase for Salmonella virulence in mice. Journal of Clinical Investigation, 95(3), 1047–1053. 10.1172/JCI117750

55. Liao, H., Zhong, X., Xu, L., Ma, Q., Wang, Y., Cai, Y., & Guo, X. (2019). Quorum-sensing systems trigger catalase expression to reverse the *oxyR* deletion-mediated VBNC state in *Salmonella* Typhimurium. Research in microbiology, 170(2), 65–73. 10.1016/j.resmic.2018.10.004

56. Romsang, A., Atichartpongkul, S., Trinachartvanit, W., Vattanaviboon, P., & Mongkolsuk, S. (2013). Gene expression and physiological role of Pseudomonas aeruginosa methionine sulfoxide reductases during oxidative stress. Journal of Bacteriology, 195(15), 3299–3308. 10.1128/JB.00167-13

57. 57. Zhao, C., Hartke, A., La Sorda, M., Posteraro, B., Laplace, J. M., Auffray, Y., & Sanguinetti, M. (2010). Role of methionine sulfoxide reductases A and B of Enterococcus faecalis in oxidative stress and virulence. Infection and Immunity, 78(9), 3889–3897. 10.1128/IAI.00165-10

58. Singh, V. K., Vaish, M., Johansson, T. R., Baum, K. R., Ring, R. P., Singh, S., Shukla, S. K., & Moskovitz, J. (2015). Significance of four methionine sulfoxide reductases in Staphylococcus aureus. PLoS ONE, 10(2), 1–20. 10.1371/journal.pone.0117594

59. Christman, M. F., Morgan, R. W., Jacobson, F. S., & Ames, B. N. (1985). Oxidative Stress and Some Heat-Shock Proteins in *Salmonella* typhimurium. Cell, 41(3), 753– 762.

60. Storz, G., Tartaglia, L. A., & Ames, B. N. (1990). Transcriptional regulator of oxidative stress-inducible genes: Direct activation by oxidation. Science, 248(4952), 189–194. 10.1126/science.2183352

61. Zheng, M., & Storz, G. (1998). Activation of the OxyR Transcription Factor by Reversible Disulfide Bond Formation. Science, 279(5357), 1718–1722.

62. Lee, C., Lee, S. M., Mukhopadhyay, P., Kim, S. J., Lee, S. C., Ahn, W. S., Yu, M. H., Storz, G., & Ryu, S. E. (2004). Redox regulation of OxyR requires specific disulfide bond formation involving a rapid kinetic reaction path. Nature Structural and Molecular Biology, 11(12), 1179–1185. 10.1038/nsmb856

63. Pomposiello, P. J., & Demple, B. 2001. Redox-operated genetic switches: the SoxR and OxyR transcription factors. Trends in biotechnology, 19(3), 109–114. 10.1016/s0167-7799(00)01542-0

64. Komitopoulou, E., Bainton, N. J., & Adams, M. R. (2004). Oxidation-reduction potential regulates RpoS levels in *Salmonella* Typhimurium. Journal of Applied Microbiology, 96(2), 271–278. 10.1046/j.1365-2672.2003.02152.x

65. Jones, D. P., & Go, Y. M. (2010). Redox compartmentalization and cellular stress. *Diabetes*, Obesity and Metabolism, 12(*SUPPL. 2*), 116–125. 10.1111/j.1463-1326.2010.01266.x

66. Dhandayuthapani, S., Jagannath, C., Nino, C., Saikolappan, S., & Sasindran, S. J. (2009). Methionine sulfoxide reductase B (MsrB) of *Mycobacterium smegmatis* plays a limited role in resisting oxidative stress. *Tuberculosis (Edinburgh*, Scotland), 89 Suppl 1(Suppl 1), S26–S32. 10.1016/S1472-9792(09)70008-3

67. Lai, L., Sun, J., Tarafdar, S., Liu, C., Murphy, E., Kim, G., & Levine, R. L. (2019). Loss of methionine sulfoxide reductases increases resistance to oxidative stress. Free Radical Biology and Medicine, 145(October), 374–384. 10.1016/j.freeradbiomed.2019.10.006

68. Trujillo-Hernandez, J. A., & Levine, R. L. (2023). Response to oxidative stress of AML12 hepatocyte cells with knockout of methionine sulfoxide reductases. Free Radical Biology and Medicine, 205(March), 100–106. 10.1016/j.freeradbiomed.2023.05.028

69. Zhang. Y., Beate Heym, Bryan Allen, D. Y. S. C. (1992). The Catalase-Peroxidase Gene and Isoniazid Resistance of Myc. Nature, 358, 591–593.

70. Hazbón, M. H., Brimacombe, M., Del Valle, M. B., Cavatore, M., Guerrero, M. I., Varma-Basil, M., Billman-Jacobe, H., Lavender, C., Fyfe, J., García-García, L., León, C. I., Bose, M., Chaves, F., Murray, M., Eisenach, K. D., Sifuentes-Osornio, J., Cave, M. D., De León, A. P., & Alland, D. (2006). Population genetics study of isoniazid resistance mutations and evolution of multidrug-resistant *Mycobacterium tuberculosis*. Antimicrobial Agents and Chemotherapy, 50(8), 2640–2649. 10.1128/AAC.00112-06

71. 71. Loewen, P. C., De Silva, P. M., Donald, L. J., Switala, J., Villanueva, J., Fita, I., & Kumar, A. (2018). KatG-Mediated Oxidation Leading to Reduced Susceptibility of Bacteria to Kanamycin. ACS Omega, 3(4), 4213–4219. 10.1021/acsomega.8b00356

72. Pesingi, P. K., Kumawat, M., Behera, P., Dixit, S. K., Agarwal, R. K., Goswami, T. K., & Mahawar, M. (2017). Protein-L-isoaspartyl methyltransferase (PIMT) is required for survival of *Salmonella* Typhimurium at 42°C and contributes to the virulence in poultry. Frontiers in Microbiology, 8(MAR), 1–9. 10.3389/fmicb.2017.00361

73. Datsenko, K. A., & Wanner, B. L. (2000). One-step inactivation of chromosomal genes in Escherichia coli K-12 using PCR products. Proceedings of the National Academy of Sciences of the United States of America, 97(12), 6640–6645. 10.1073/pnas.120163297

74. Aebi, H. (1984). Catalase in Vitro. Methods in Enzymology, 105(C), 121–126. 10.1016/S0076-6879(84)05016-3

75. Noble, R. W., & Gibson, Q. H. (1970). The reaction of ferrous horseradish peroxidase with hydrogen peroxide. Journal of Biological Chemistry, 245(9), 2409–2413. 10.1016/s0021-9258(18)63167-9

76. Chomczynski, P., & Sacchi, N. (1987). Single-step method of RNA isolation by acid guanidinium thiocyanate-phenol-chloroform extraction. Analytical Biochemistry, 162(1), 156–159. 10.1016/0003-2697(87)90021-2

77. Livak, K. J., & Schmittgen, T. D. (2001). Analysis of relative gene expression data using real-time quantitative PCR and the 2(-Delta Delta C(T)) Method. *Methods (San Diego*, Calif.), 25(4), 402–408. 10.1006/meth.2001.1262

78. Ellman, G. L. (1959). Tissue sulfhydryl groups. Archives of biochemistry and biophysics, 82(1), 70–77. 10.1016/0003-9861(59)90090-6

79. Eyer, P., Worek, F., Kiderlen, D., Sinko, G., Stuglin, A., Simeon-Rudolf, V., & Reiner, E. (2003). Molar absorption coefficients for the reduced ellman reagent: Reassessment. Analytical Biochemistry, 312(2), 224–227. 10.1016/S0003-2697(02)00506-7

