## supplemental tables for "Compensatory role of KatG in defending H_2_O_2_ stress in *msr* deletion strain of *Salmonella* Typhimurium"

**Supplementary Table S1: Primer sequence for real time PCR**

| Gene | Primer sequence | References |
|---|---|---|
| *gmk*<br>Forward Primer<br>Reverse Primer | <br>5’–TTGGCAGGGAGGCGTTT–3’<br>5’–GCGCGAAGTGCCGTAGTAAT–3’ | Nikaido *et al.,* 2012 |
| *katE*<br>Forward primer<br>Reverse primer | <br>5’–ATTCCGGAAGAGTTGGTGCCAGTA–3’<br>5’–GACGACTGATTTGCGTGTCGGTAT–3’ | Kirthika *et al.,* 2022 |
| *katG*<br>Forward primer<br>Reverse primer | <br>5’–TTAACTCCTGGCCGGATAAC–3’<br>5’–TAATCGGCCACAACAAACG–3’ | Kirthika *et al.,* 2022 |
| *katN*<br>Forward primer<br>Reverse primer | <br>5’–GCGCGAGCGAAGATCATTTAT–3’<br>5’–GCGACTTCACGGGTCATTAAGA–3’ | Kirthika *et al.,* 2022 |

**Supplementary Table S2: Primer sequence for *katG* deletion**

| | Primer name | Sequence | Amplicon size |
|---|---|---|---|
| 1 | *katG* deletion forward | 5’-AGCTCCCTGCCGGGAGCTTTATTACAACT CATATTGATCTGTAGGCTGGAGCTGCTTC-3’ | 1092 bp |
| 2 | *katG* deletion reverse | 5’- TTATTGCAGATCGAAACGGTCCAGGTTCA TCACTTTCACCCATATGAATATCCTCCTTA-3’ | |
| 3 | *katG* deletion test forward | 5’- TCTGCCGGTATCGACGTAGA-3’ | 1607 bp |
| 4 | *katG* deletion test reverse | 5’- CCTGGTTCCAGCGTCAAAAC-3’ | |
